# mRNA display in cell lysates enables identification of cyclic peptides targeting the BRD3 extraterminal domain

**DOI:** 10.1101/2024.03.20.585962

**Authors:** Catherine A. Hurd, Jacob T. Bush, Andrew J. Powell, Louise J. Walport

## Abstract

mRNA display is a powerful technology to screen libraries of >10^12^ cyclic peptides against a protein target, enabling the rapid discovery of high affinity ligands. These cyclic peptides are particularly well suited to challenging protein targets that have been difficult to drug with small molecules. However, target choice can still be limited as screens are typically performed against purified proteins which often demands the use of isolated domains and precludes the use of aggregation-prone targets. Here, we report a method to perform mRNA display selections in mammalian cell lysates without the need for prior target purification, vastly expanding the potential target scope of mRNA display. We have applied the methodology to identify low to sub-nanomolar peptide binders for two targets; a NanoLuc subunit (LgBiT) and full-length bromodomain-containing protein 3 (BRD3). Our cyclic peptides for BRD3 were found to bind to the extraterminal (ET) domain of BRD3 and the closely related BRD proteins, BRD2 and BRD4. While many chemical probes exist for the bromodomains of BRD proteins, the ET domain is relatively underexplored, making these peptides valuable additions to the BRD toolbox.

## Introduction

Macrocyclic peptides are a promising class of potential therapeutics, offering exquisite selectivity and tight binding affinities in addition to high proteolytic stability.^1^ Their increased interaction area relative to small molecules allow them to make extensive contacts with relatively featureless surfaces, such as those involved in protein-protein interactions (PPIs). This property has enabled peptide inhibitors to be identified for a broad range of targets previously thought of as ‘undruggable’.^2,3^ In addition to their use as therapeutics, cyclic peptides make attractive chemical tools for target validation, interrogating biological pathways and as pulldown agents to study protein interactions.^4–6^

The rise in popularity of macrocyclic peptides has been facilitated by the development of *in vitro* display technologies for their discovery. These technologies include phage display and mRNA display, both of which can be used to rapidly identify high-affinity hits from libraries with billions to trillions of starting members.^7–12^ The RaPID (Random non-standard peptide integrated discovery) system combines flexible *in vitro* translation (FIT) with mRNA display to enable the generation of covalently-cyclized peptide libraries containing non-canonical amino acids.^13–15^ Peptide display technologies have been successful in identifying high-affinity ligands for a broad range of targets, including several recent examples that have progressed to late stage clinical trials.^16,17^

Currently, the affinity-panning step used to identify binders from peptide display libraries requires an immobilized protein target, which typically demands access to highly purified, heterologously expressed protein. This is incompatible with aggregation-prone, intrinsically disordered or membrane-bound targets, limiting target scope. When screening DNA-encoded small molecule libraries (DELs), where a similar affinity-panning step is used, several groups have pioneered strategies to circumvent the need for recombinant protein by screening these libraries directly in lysates,^18,19^ inside living cells or on the cell surface.^20–22^ Similar approaches have not yet been explored for mRNA-display libraries. The ability to perform RaPID selections in cell lysates would increase the scope of possible target proteins to include both proteins that are challenging to express and purify, and also those requiring post-translational modifications or native binding partners/cofactors to retain their active form.

Here, we report a method to perform RaPID selections against target proteins expressed in mammalian cell lysates. We apply this method to identify low to sub-nanomolar cyclic peptide binders of both a NanoLuc subunit (LgBiT) and full length bromodomain-containing protein 3 (BRD3). Our new method will increase the scope of targets that can be screened by mRNA display, enable selections to be performed in more biologically relevant environments and remove the time-consuming process of producing purified proteins in their active forms.

## Results and Discussion

We sought to apply the RaPID methodology to targets expressed in mammalian cell lysates. In the method described here, we envisaged exogenously overexpressing the target protein in mammalian cells. Following cell lysis, the lysate would be incubated with the mRNA-peptide library followed by antibody-coated magnetic beads to immunoprecipitate the target protein and any peptides bound to it. The remaining lysate proteins and any non-bound peptides would then be removed by washing the beads and hit sequences eluted from the beads by boiling and amplified by PCR for further selection rounds (Figure 1).

**Figure 1.**
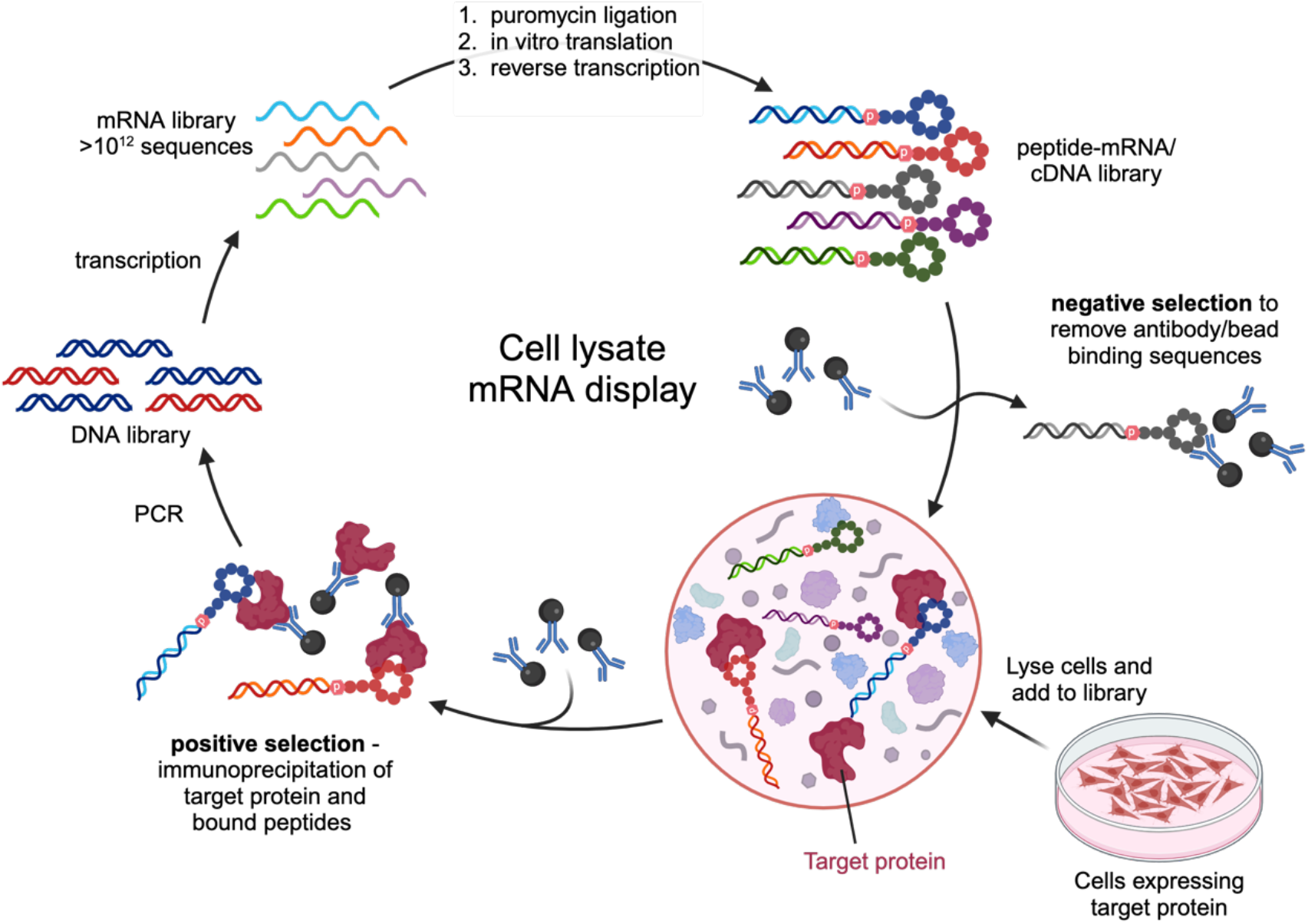
Cell lysate mRNA display method. Transcription, puromycin ligation, translation and reverse transcription are performed as in a typical RaPID selection.^23^ The library is then panned against antibody-coated beads to remove sequences binding to either the antibody or the beads. This depleted library is then added to cell lysate expressing a target protein and the target is immunoprecipitated using antibody-coated beads. Following washes to remove lysate proteins and non-selected peptides, the DNA barcodes from positive hits are amplified by PCR to carry into the next round of selection.

Before undertaking selections with target proteins in cell lysates, we sought to address the following potential issues: (1) whether sufficient expression levels of the target protein could be achieved, (2) whether the protein could be efficiently captured via antibody immobilization and (3) whether the peptide-encoding RNA/DNA hybrid would be degraded in a lysate. The NanoBiT system,^24^ was chosen as an ideal test system as the concentration of LgBiT protein in cell lysates can be accurately measured using luminescence assays through complementation with a known high-affinity peptide binder, HiBiT (*K*_D_ = 0.7 nM,^24,25^ Figure 2A). We first prepared a 3xFlag-tagged LgBiT construct for mammalian expression. Following transfection of HEK 293T cells with increasing quantities of 3xFlag-LgBiT, the protein was quantified in luminescence assays through complementation with excess HiBiT (Figure S1). LgBiT concentrations up to 1 µM could be reached, exceeding our typical selection conditions of 200 nM, and the protein could be efficiently immunoprecipitated with anti-Flag antibody-coated beads (Figure S2). To assess the stability of our mRNA libraries, a reverse-transcribed mRNA library was incubated in a HEK 293T cell lysate and the DNA quantity monitored over time by qPCR (Figure S3). No loss of DNA was observed indicating that the DNA/RNA hybrid is not degraded under the proposed selection conditions.

**Figure 2.**
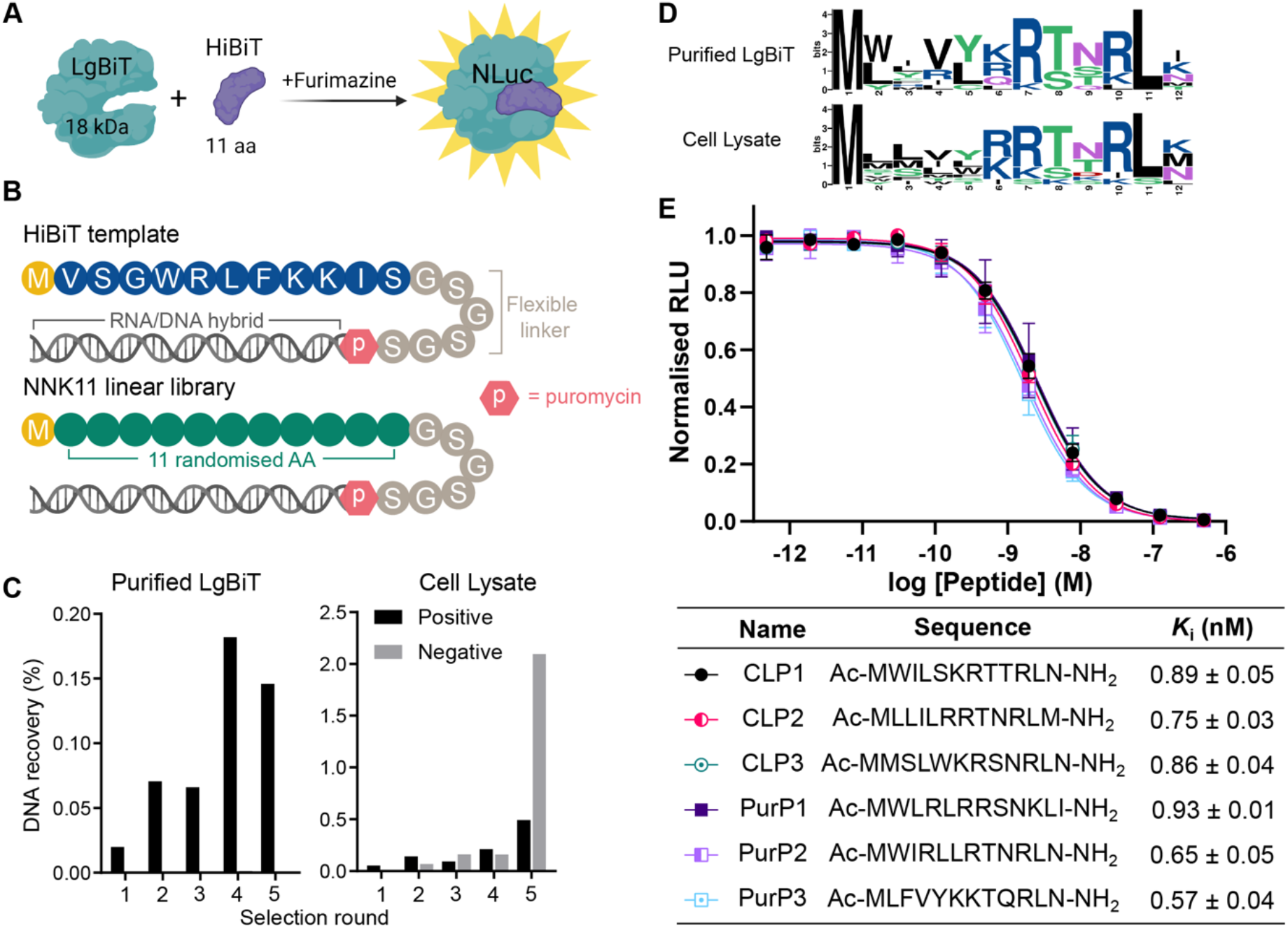
LgBiT test selections. **A**. NanoBiT® system from Promega. The HiBiT peptide complements the LgBiT protein to produce functional nanoluciferase (NLuc). **B**. Library design for LgBiT selections. NNK codons (N = A/T/C/G, K = T/G) in the NNK11 linear library can code for any of the 20 canonical amino acids. The peptide sequences are followed by a flexible GSGSGS linker. **C**. DNA recoveries from positive and negative selections at the end of each round as a percentage of the input DNA library. **D**. Logo plot analysis of the top ten sequences identified from each selection, generated using the Berkeley WebLogo server.^26^ **E**. Displacement of HiBiT-HaloTag control protein (24 nM) by the top three peptides identified from cell lysate (CLP1-3) and purified protein (PurP1-3) RaPID selections in luminescence competition assays. Curves and calculated *K*_i_ values are given as the mean ± SEM (calculated from at least three independent replicates). Ac – acetylated N-terminus.

Having confirmed that sufficient target protein expression and capture could be achieved, and that the library was stable under the desired selection conditions, we performed a full RaPID selection using the LgBiT protein as a model target. The selection was performed against 3xFlag-tagged LgBiT overexpressed in crude HEK 293T cell lysates, using anti-Flag antibody-coated beads for immunoprecipitation. For comparison we also performed a selection against biotinylated LgBiT that had been purified from *E. coli* and immobilised on streptavidin beads, representing typical selection conditions. We anticipated that enrichment of similar hit sequences from both selections would demonstrate that the cell lysate method can identify *de novo* peptide ligands from a randomised library in a similar manner to the well-validated methodology using purified proteins. An mRNA library was prepared, encoding linear peptides containing a Met initiator codon followed by 11 randomised amino acids to mimic the length of the known high-affinity HiBiT peptide (Figure 2B). The starting diversity of this library was estimated to be >10^12^ unique sequences. Approximately 100 copies of the HiBiT sequence were also spiked into the library as a positive control (Figure 2B); round-by-round enrichment of the HiBiT sequence would confirm that the selection was working as intended.

Following a standard affinity panning protocol, during each round of selection, negative selections were first performed in which translated libraries were incubated with selection beads lacking the protein of interest (i.e. streptavidin beads or anti-flag antibody-coated beads) to deplete bead-binding sequences (Figure 1). Translated libraries now depleted of bead-binding sequences were then incubated with target protein in a ‘positive selection’. While sequences captured in both negative and positive selections were retained for quantification and downstream sequencing, only sequences captured in the positive selection were carried through to the next selection round. Negative selection recoveries are expected to remain low throughout the selection rounds while the positive selection recoveries increase. However, while this occurred in the selection with purified protein, in the cell lysate selection against Flag-tagged LgBiT, negative selection recoveries increased throughout the selection (Figure 2C). Nonetheless, following five rounds of selection, libraries were sent for next generation sequencing (NGS). Analysis of the NGS data revealed that the high negative recoveries in the cell lysate selection were due to enrichment of Flag-like sequences containing DYKxxD motifs (Supplementary File 1,2). This suggested that more exhaustive negative selections are needed to deplete bead-binding sequences when using antibody immunoprecipitation. Despite this high background of antibody-binding sequences, 69 of the top 100 most enriched sequences from the Round 5 positive selection appeared at much higher frequency in the positive selection sequencing than in the negative selection sequencing (at least 30-fold). This suggested that peptide sequences specifically binding to the LgBiT protein had been successfully enriched. We were surprised to find that the spiked HiBiT sequence had not been enriched during either selection, though we could confirm its presence in our starting libraries as the sequence was observed at a low frequency in the early selection rounds.

To compare the hits from each selection, logo plots were generated using the top 10 most enriched sequences identified in the Round 5 positive sequencing data (Figure 2D). An almost identical consensus sequence was revealed in both selections, comprising hydrophobic residues at the N-terminus and a highly conserved (K/R)RTNRL motif towards the C-terminus. To confirm the binding of the hit peptides to LgBiT protein, the top 3 hits from the purified selection (PurP1-3) and cell lysate selection (CLP1-3) were produced using solid phase peptide synthesis (Table S1). In addition, we synthesised the HiBiT peptide with an N-terminal methionine as was spiked into the selections. This M_HiBiT peptide produced functional NLuc upon binding to LgBiT with an apparent *K*_D_ of 2.7 ± 0.2 nM (Figure S4), which is in line with the previously reported affinity for the HiBiT peptide (0.7 nM).^25^ The peptide hits identified in our selections did not produce functional NLuc upon incubation with LgBiT, however these peptides were able to displace a HiBiT-tagged HaloTag protein in luminescence competition assays, allowing for their *K*_i_ values to be determined (Figure 2E). This demonstrated that the peptides share the same binding site on LgBiT as the HiBiT peptide despite not generating a productive luciferase. This has been shown previously with the DrkBiT peptide, where a single Arg to Ala substitution from the HiBiT sequence ablates luminescence activity.^27^ The hit peptides from both of our selections showed high-affinity binding to LgBiT with *K*_i_ values ranging from 0.5 to 0.9 nM, exceeding that of the HiBiT peptide and validating that our cell lysate selection method is suitable for the identification of novel peptide ligands.

We next sought to apply our cell lysate selection methodology to a more challenging target, Bromodomain-containing protein 3 (BRD3). The full-length protein contains several unstructured regions and has not been amenable to purification for screening campaigns using established methods (Figure 3A).^28^ This presented an opportunity for our lysate-based methodology to select peptides against the full-length protein in its biologically relevant conformation, and could lead to the identification of ligands engaging multiple regions of the protein. Over the past decade, there has been substantial effort to develop small molecule inhibitors of the BRD proteins as anti-cancer agents.^29–32^ To date, all molecules that have entered clinical trials target the bromodomains, BD1 and BD2. A recent study, however, identified fragments that weakly engage the extraterminal (ET) domain.^33^ mRNA display selections have also been used to identify high-affinity cyclic peptides for the individual bromodomains.^34,35^ Before embarking on a RaPID selection, we first confirmed that the full-length protein could be expressed and immunoprecipitated from HEK 293T cells in sufficient quantities to recover a previously discovered cyclic peptide in a mock selection round (Figure S5). For this, we used a HiBiT-tagged BRD3 construct, enabling both quantitation in lysates using luminescence assays and protein immobilization on magnetic beads via an anti-HiBiT antibody. In contrast to LgBiT, much lower concentrations of full-length BRD3 were present in the lysate following transfection (∼40 nM), and we therefore decided to immunoprecipitate the protein and directly resuspend this bead-bound protein in lysate at approximately 200 nM immediately prior to the positive selection step.

**Figure 3.**
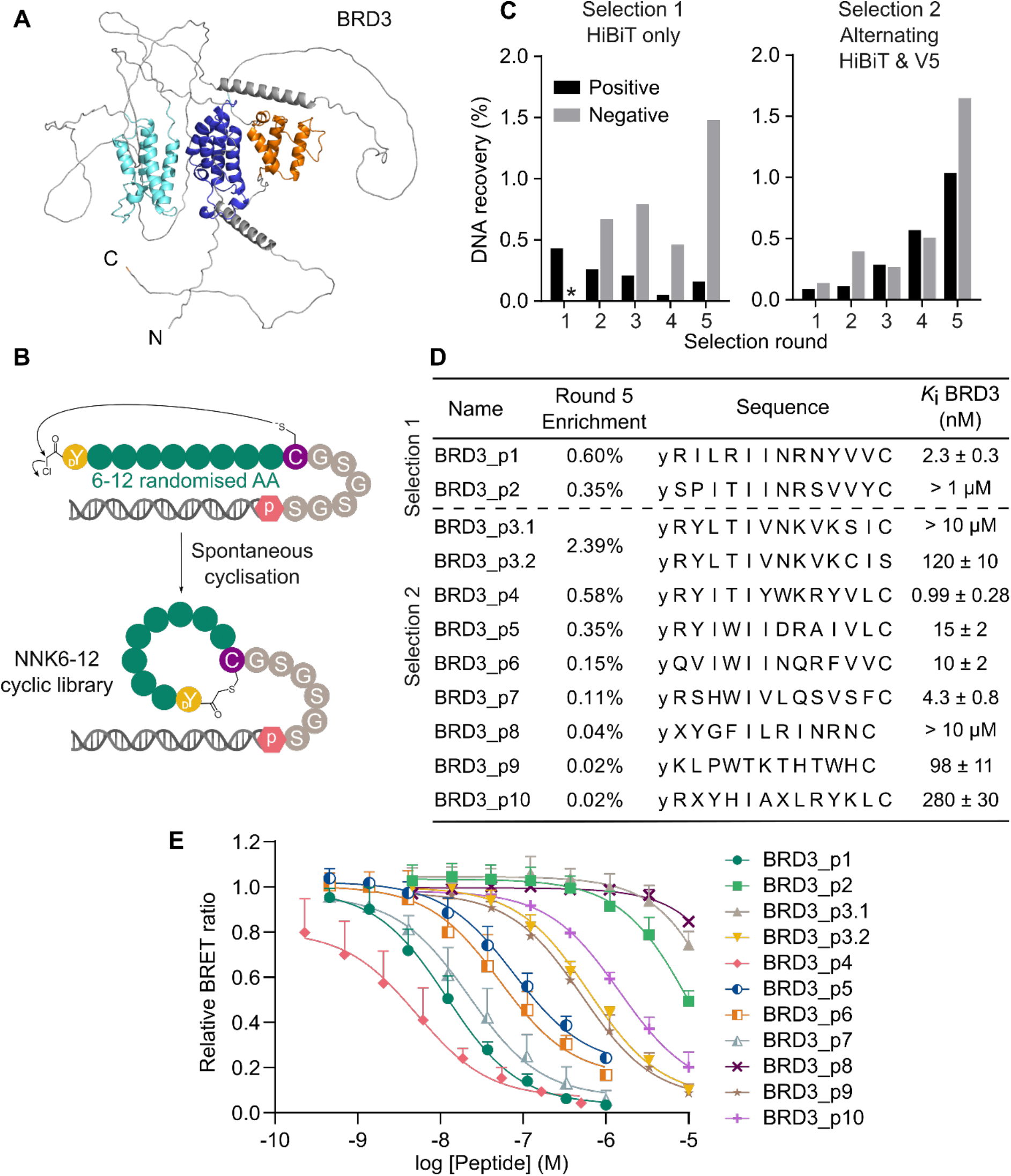
Cell lysate selections against full-length BRD3. **A**. Alphafold predicted structure of BRD3 with bromodomains (cyan and blue) and ET domain (orange) coloured as in Fig. 4A. **B**. BRD3 library design. Peptides were initiated with ClAc-_D_Tyr followed by 6-12 randomised amino acids (AA) and a cysteine for cyclisation with the N-terminal chloroacetyl group to produce a thioether-cyclised library. **C**. DNA recoveries throughout the RaPID selections against full-length BRD3. In the HiBiT only selection, no negative selections were performed in Round 1 (denoted ^*^). **D**. Round 5 NGS enrichments of selected sequences identified from RaPID selections against full-length BRD3. y = _D_Tyr and X = *N*-methylalanine. BRD3_p1 and BRD3_p2 were identified in Selection 1, while BRD3_p3 to BRD3_p10 were identified in Selection 2. Two peptides were synthesised based on the BRD3_p3 sequence due to the occurrence of a Cys in the randomised region giving rise to two possible cyclisation points. In each peptide the Cys not used for cyclisation was replaced with a Ser for ease of synthesis. *K*_i_ values for BRD3 are calculated from the NanoBRET curves in E. Values are given as the mean ± one standard error of the mean (calculated from at least three independent replicates). **E**. Displacement of BRD3_p6_TAMRA (100 nM) from BRD3 by peptides in NanoBRET competition assays. Values are plotted as the mean + one standard error of the mean (calculated from at least three independent replicates).

Having confirmed the suitability of our construct in the mock selection, we prepared mRNA libraries encoding cyclic peptides with 6-12 variable amino acids within the macrocycle (Figure 3B). Flexizyme-mediated genetic code reprogramming was used to initiate all peptides with *N*-chloroacetyl-_D_-tyrosine (ClAc-_D_Y) to allow for macrocyclization with a down-stream cysteine residue as described previously,^36^ and AUG (Met) codons appearing within the macrocycle were reprogrammed to incorporate *N*-methyl-alanine. Using the protocol established in the mock selection, the cyclic peptide library was panned against HiBiT-tagged full-length BRD3 over 5 rounds of selection and recovered DNA was analysed by NGS (Figure 3C). To combat the enrichment of bead-binding sequences observed in the LgBIT selection, we initially increased our negative selection stringency to include five negative selections in every round from Round 2 onwards. Despite these stringent negative selections, high negative recoveries were observed and NGS analysis revealed that HiBiT-like motifs were significantly enriched during the selection; more than 950 of the top 1000 sequences in the positive selection contained motifs that were also significantly enriched in antibody-only negative selections (GWR/K or GWxW – Supplementary File 3), suggesting that these peptides are binding directly to the HiBiT antibody rather than BRD3. However, even with this high background, several sequences identified were only enriched in the positive selection, suggesting these might be binding to the BRD3 protein. Alignment of these sequences revealed a consensus sequence with strong preference for Arg or Leu in position 2, a highly conserved Ile at position 6, and a preference for cationic Arg or Lys in positions 8 and 9 (Figure S6).

To enable more target-binding hits to be identified during the selection, we aimed to further reduce the high recoveries of sequences binding to the immobilised antibody. To this end, we decided to swap the antibody and tags used for immuno-precipitation between selection rounds. As an alternative to the HiBiT tag, HEK 293T cells were transfected with a V5-tagged full-length BRD3 construct and a V5 antibody was used for immunoprecipitation (Figure S7). Additionally, we performed extensive negative selections with the translated library in Round 1, a step that is often excluded to preserve library diversity. The selection was repeated using immunoprecipitation via a HiBiT tag in rounds 1, 3 & 5, and a V5 tag in rounds 2 & 4 (Figure 3C). Pleasingly, while the negative selection recoveries were still higher than in a standard RaPID screen, good positive recoveries were observed and NGS analysis showed that no enrichment of antibody-binding motifs occurred in either the negative or positive selections; the GWR/K or GWxW motifs were no longer observed in the top 1000 sequences from either selection. Instead, we observed enrichment of many related sequence families in the positive selections only, indicating that these sequences are specifically binding to BRD3 (Supplementary file 4) Alignment of the most enriched sequences showed a very similar consensus sequence to the first selection, with Ile at position 6 in almost all sequences, and a strong preference for Arg residues at positions 2 and 9. Val and Ile were the most commonly observed residues at positions 12 and 13 in both selections (Figure S6).

As we are proposing a method that enables screening of non-purified proteins in lysates, we also wanted to develop methods to validate the peptide hits without the need for target purification, as is demanded by many standard biophysical assays. We selected 11 peptide sequences for solid phase synthesis to validate their binding to BRD3 (Figure 3D). Peptides were selected to cover a range of enrichment values and to include diverse sequences that could be binding to different regions on BRD3. The peptides were synthesised with an azide group at the C-terminus that could be functionalized to enable the use of a range of assays (Table S2). In the first instance, several peptides were modified with a TAMRA fluorophore to monitor direct binding to the HiBiT-tagged full-length BRD3 in HEK 293T lysates using NanoBRET assays (Table S3).^37^ Peptides BRD3_p5_TAMRA and BRD3_p6_TAMRA were found to bind to full-length BRD3 with *K*_D_ values of 21 ± 3 nM and 23 ± 4 nM, respectively (Figure S8). The remaining peptides were tested in NanoBRET competition assays, displacing BRD3_p6_TAMRA, to assess whether peptides bind to the same site on BRD3 and to measure their *K*_i_ values where this was the case (Figure 3C-E). Eight out of 11 of the peptides tested fully competed with BRD3_p6_TAMRA, demonstrating that they share a common binding site. Two of the peptides that only partially competed with BRD3_p6_TAMRA (BRD3_p2 and BRD3_p8) were themselves labelled with TAMRA and tested in direct binding assays to assess whether the peptides were binding to a different site on BRD3. However, both showed only modest affinity for direct binding to the full-length protein, in agreement with the weak competition that was observed, rather than tight binding at an alternate binding site (Figure S8).

To identify the binding site of the peptides, HiBiT-tagged truncated constructs of BRD3 were prepared. These included the N-terminal region encompassing the two bromodomains (1-428), the C-terminal portion containing the ET domain (429-726) and the full-length construct with the ET domain deleted (ΔET) (Figure 4A). The binding of BRD3_p5_TAMRA to these constructs was measured using direct NanoBRET assays; binding was preserved with BRD3(429-726) but no binding to the bromodomains or the construct lacking the ET domain (ΔET) was observed (Figure 4A). This indicated that the BRD3 ET domain is required for peptide binding. To investigate this further, two constructs were expressed and purified from *E. coli* as biotinylated proteins; The BRD3-L construct, encompassing residues 420-726 including the ET domain and surrounding unstructured regions, and the BRD3-ET construct (562-644), truncated at the boundaries of the ET domain. Binding of the peptides to these constructs was measured using surface plasmon resonance (SPR) (Table 1, Figure S9 and S10). All peptides bound to the two constructs with very similar affinities, indicating that the primary binding site of the peptides is the ET domain and that the surrounding residues are not forming significant interactions with the peptides. The *K*_D_ values determined for the ET domain were in close agreement with the *K*_i_ values measured for the full-length BRD3 in NanoBRET assays, however the ranking of the peptides varied slightly between these experiments. In particular, BRD3_p4 and BRD3_p7 appeared to bind more tightly to the full-length protein in NanoBRET than to the isolated ET domain in SPR experiments. This could have been due to additional interactions with the bromodomains in the full-length construct, therefore we tested their binding to the truncated BRD3(429-726) construct lacking the bromodomains in NanoBRET competition assays (Figure S11). Both peptides bound this construct with a very similar affinity to the full-length protein, demonstrating that the bromodomains are not involved in binding. Based on these experiments, it appears that all peptides that were tested bind solely to the ET domain and that the minor observed differences between NanoBRET *K*_i_ values and SPR *K*_D_ values are due to assay variations.

**Table 1:** Calculated *K*_D_ values from SPR experiments. Values are given as the mean ± one standard deviation from two replicates.

| | $K_D$ BRD3-ET (NM) | $K_D$ BRD3-L (NM) |
| --- | --- | --- |
| BRD3_P1 | $4.7 \pm 0.3$ | $3.3 \pm 0.4$ |
| BRD3_P2 | $> 1 \mu\text{M}$ | $> 1 \mu\text{M}$ |
| BRD3_P3.1 | $> 1 \mu\text{M}$ | $680 \pm 280$ |
| BRD3_P3.2 | $45 \pm 2$ | $51 \pm 3$ |
| BRD3_P4 | $8.0 \pm 0.7$ | $5.9 \pm 0.5$ |
| BRD3_P5 | $13 \pm 1$ | $13 \pm 4$ |
| BRD3_P6 | $4.8 \pm 0.7$ | $4.2 \pm 0.3$ |
| BRD3_P7 | $21 \pm 2$ | $17 \pm 2$ |
| BRD3_P8 | $> 1 \mu\text{M}$ | $> 1 \mu\text{M}$ |
| BRD3_P9 | $28 \pm 1$ | $38 \pm 2$ |
| BRD3_P10 | $110 \pm 10$ | $250 \pm 40$ |

**Figure 4.**
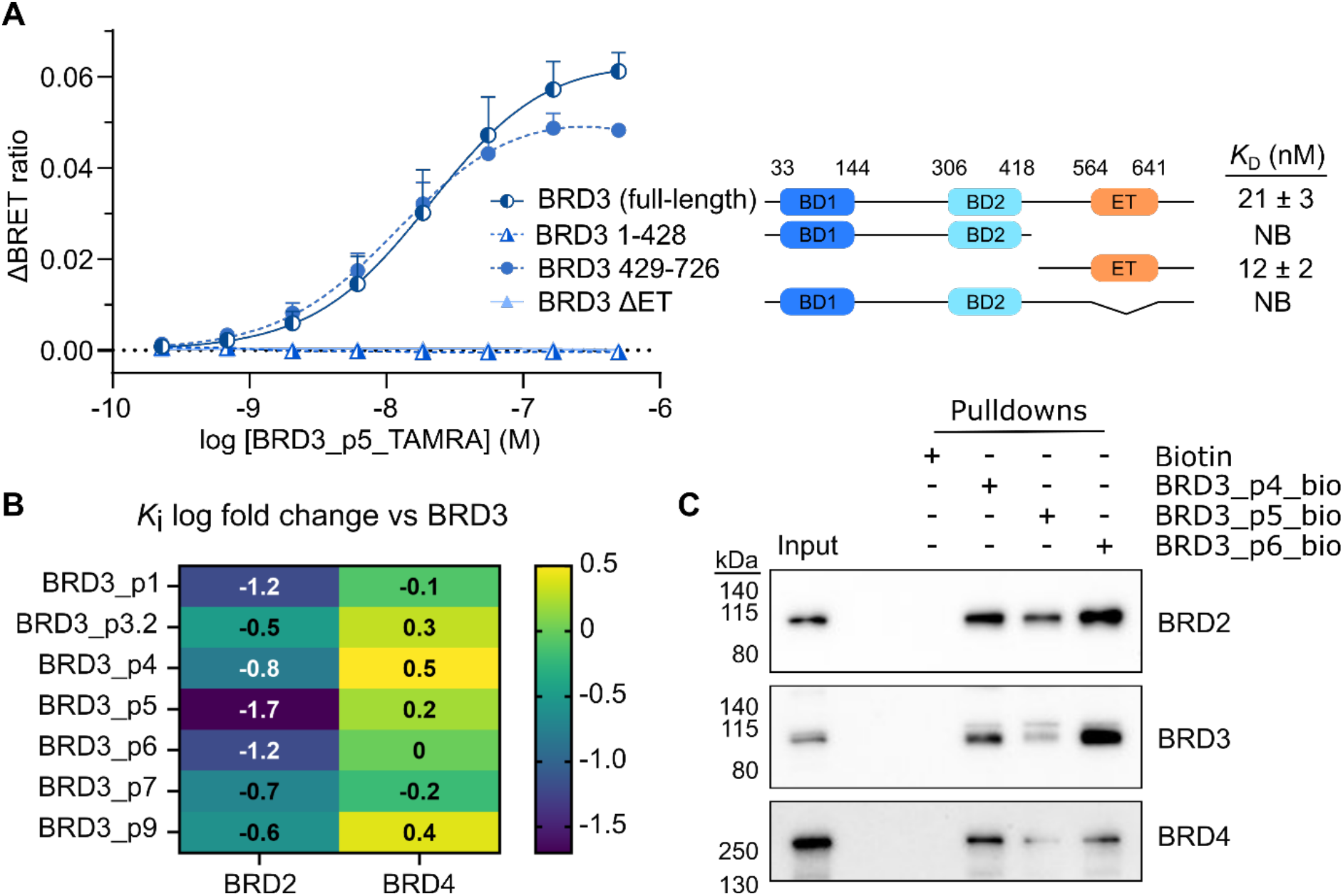
Peptide interactions with ET domain and other BRD proteins. **A**. Direct binding of BRD3_p5_TAMRA to HiBiT-tagged BRD3 constructs in NanoBRET assays. **B**. *K*_i_ log fold change between peptides binding to BRD2 and BRD4, compared with BRD3. NanoBRET curves and *K*_i_ values are shown in Figure S12. **C**. Detection of endogenous BRD proteins following biotin or biotinylated peptide pulldowns from HEK 293T nuclear extracts. Pulldown samples are concentrated ten-fold compared to the input sample.

The ET domain is highly conserved between BRD proteins, with >80% sequence identity between BRD2, BRD3 and BRD4. Several interaction partners for the BRD ET domains have been identified, including components of the NuRD and BAF complexes involved in chromatin remodelling,^38,39^ and viral integrases.^40^ We investigated the binding of selected peptides to HiBiT-tagged full-length BRD2 and BRD4 in NanoBRET competition assays (Figure 4B and S12). All peptides tested displayed nanomolar affinities for BRD4, although there were some differences in affinity compared with BRD3 e.g. BRD3_p4 and BRD3_p9 showed *K*_i_ values that were three-fold tighter to BRD4, while BRD3_p1 and BRD3_p6 bound with near identical affinities for the two proteins. The observed binding to BRD2 was weaker for all peptides tested, with differences ranging from three-fold to > 50-fold. This was surprising given the high levels of sequence identity between the BRD ET domains, but likely reflects the common observation that cyclic peptides exploit multiple interactions across a binding surface such that only small amino acid changes or changes in dynamics can alter peptide binding.^34,41,42^ It has been shown previously that the ET domain of BRD3 and BRD4 engage binding partners with a common ‘KIKL’ motif, and linear peptides derived from the interaction partners have been shown to bind the BRD3 ET domain with micromolar affinities.^38^ Some of our hits contain similarities to these sequences, such as the KVK motif in our top hit (BRD3_p3), however the affinities of our cyclic peptides for the BRD3 ET domain greatly exceeds the linear peptides tested previously, suggesting they may be using an alternative binding mode.

During our selection, we used a tagged form of BRD3 that had been overexpressed in HEK 293T cells. As such, we wanted to confirm that the peptides would also engage the endogenous protein using pulldowns. Peptides BRD3_p4, BRD3_p5 and BRD3_p6 were biotinylated via a click reaction with biotin-alkyne, and the biotinylated peptides were immobilized on streptavidin beads and incubated with HEK 293T nuclear extracts. Western blotting revealed that endogenous BRD2, BRD3 and BRD4 were detected in the pulldown samples for all three peptides, while no pulldown was seen with biotin alone (Figure 4C). This demonstrated that the peptides can interact with the endogenous BRD proteins and be used as efficient pulldown probes.

## Conclusions

Here we have reported a method in which full length protein targets can be screened by mRNA display in crude cell lysates without the need for prior purification. We have demonstrated our selection method with LgBiT and BRD3 proteins in HEK 293T cell lysates. Both selections were successful in identifying novel peptide ligands from a starting library of >10^12^ mRNA sequences, and the hit peptides displayed affinities in the sub- to low-nanomolar range when tested in subsequent assays.

We envisage our method will be applicable to a diverse range of protein targets, enabling discovery of cyclic peptide binders of proteins in their full-length form and native conformations. In addition to widening target scope, screening proteins in crude lysate will save the substantial time required to generate purified, active protein constructs, which can be particularly challenging for aggregation-prone and intrinsically disordered proteins, even in instances where it is possible. In some cases, a challenge for lysate-based screening may be target concentration. Here we have overexpressed the protein of interest in mammalian cells to achieve high expression. It is possible, however, that sufficient concentrations of endogenous target protein could be achieved by immunoprecipitation immediately prior to selection, as was used for the BRD3 selections. Given the potency of peptides identified by mRNA display, it is also likely that target concentrations of much less than 200 nM could be used in future selections.

The HiBiT-tagging approach implemented in this work has multiple benefits, enabling target quantification and efficient immunoprecipitation during the selections, as well as rapid downstream assays to validate selection hits. In particular, the NanoBRET assay described here is a powerful method for quantifying binding affinities of peptides for proteins present at very low abundances in a crude cell lysate mixture.^37^ This assay retains sensitivity at and below endogenous protein levels. As such CRISPR/Cas9 systems could be used to introduce HiBiT tags to endogenous proteins if desired, both for use in hit validation and also potentially as a tag for immunoprecipitation if using endogenous proteins as the target in a selection.^25^ Use of the NanoBRET assay removes the need for protein purification for hit validation with standard biophysical techniques. Additionally, we demonstrated that biotinylated peptide pulldowns can be used as a complementary method to validate engagement with endogenous proteins.

Overall, we anticipate that our new screening approach will be a generally applicable and time-efficient method to perform mRNA display selections under conditions that are more biologically relevant and that give access to a broad range of targets that cannot be screened with current methodologies. This will be particularly beneficial to early-stage drug discovery projects, where the expression and purification of novel protein targets can be a challenging, time-consuming, and sometimes impossible process.

## Acknowledgements

We would like to thank the Crick Advanced Sequencing Science Technology Platform for assistance with next generation sequencing, the Crick Structural Biology Science Technology Platform for expert technical support and the Crick Cell Services Science Technology Platform for providing the cell lines used in this work. This work was supported by the Francis Crick Institute which receives its core funding from Cancer Research UK (CC2030), the UK Medical Research Council (CC2030), and the Wellcome Trust (CC2030). We also thank GSK for its commitment to support fundamental discovery research through the establishment of the Crick-GSK LinkLabs partnership, and David House for his advice and support for the project. Illustrative figures were created using BioRender.com.

