## Supplementary Data for "mRNA display in cell lysates enables identification of cyclic peptides targeting the BRD3 extraterminal domain"

### Supplementary Materials and Methods

#### S1. Materials

Unless otherwise stated reagents were purchased from commercial sources including Merck, Thermo Fisher Scientific, New England Biolabs (NEB) and Invitrogen. Primers were purchased from Integrated DNA Technologies. Primer sequences used for mRNA display are given in Table S4. Sanger sequencing was performed by Genewiz and whole plasmid sequencing by Plasmidsaurus or Full Circle.

#### S2. Cloning of expression constructs

The cDNA encoding Large bit (LgBiT, Promega),[1] BRD3-L(420-726) and BRD3-ET(562-644) was cloned into pQE80LNAviH at *Bam*HI/*Eco*RI sites. This bicistronic plasmid encodes an N-terminal Hexa-His/Avi-tag on the protein of interest and also contains the gene encoding BirA biotin ligase. Biotinylated $His_6$-Avi-tagged proteins were produced in the BL21 (DE3) *E.coli* strain (NEB).

The cDNA encoding LgBiT (Promega) was cloned into the p3xFLAG-CMV-10 vector (Sigma) at *Bam*HI/*Eco*RI sites. Full-length human BRD2, BRD3 and BRD4 (long isoform) were cloned into pcDNA-3.1D with C-terminal HiBiT tags, using vector linearisation by PCR and InFusion cloning (Takara). Truncations of BRD3 (1-428, 429-726 and ΔET(Δ564-641)) were prepared by deletion mutagenesis using pcDNA3.1D/BRD3-HiBiT as a template and Pfu Turbo polymerase (Agilent, 600250) following the manufacturer's instructions.

#### S3. Protein expression and purification

*LgBiT:* Cultures were grown in terrific broth at 37 °C to an $A_{600}$ of 0.6-0.8 before induction with 0.1 mM IPTG and 200 µM biotin for 5 h at 37 °C. Cells were harvested and lysed with 50 mM HEPES pH 7.5, 250 mM NaCl, 1 mM TCEP, 20 mM imidazole, 5% glycerol, DNase, 1 mg/mL lysozyme and cOmplete™ EDTA-free Protease Inhibitor Cocktail (Roche). Biotinylated $His_6$-Avi-LgBiT was purified at 4 °C using a HisTrap HP column (Cytiva) and eluted with 50 mM HEPES pH 7.5, 150 mM NaCl, 1 mM TCEP and 250 mM imidazole. The protein was further purified with a Superdex S75 16/600 (Cytiva) gel filtration column with 50 mM HEPES pH 7.5, 150 mM NaCl and 1 mM TCEP. Purified protein was flash frozen and stored at -80 °C.

*BRD3 constructs:* Cultures were grown in terrific broth at 37 °C to an $A_{600}$ of 0.6-0.8 before induction with 0.1 mM IPTG and 200 µM biotin overnight at 20 °C. Cells were harvested and lysed with 50 mM Tris pH 7.5, 300 mM NaCl, 1 mM DTT, 20 mM imidazole, 100 µM $MgCl_2$, DNase, 1 mg/mL lysozyme and cOmplete™ EDTA-free Protease Inhibitor Cocktail (Roche). Biotinylated $His_6$-Avi-tagged BRD3 proteins were purified at 4 °C using a HisTrap HP column (Cytiva) and eluted with 50 mM Tris pH 7.5, 150 mM NaCl, 1 mM DTT and 250 mM imidazole. The proteins were further purified with a Superdex S75 16/600 (Cytiva) gel filtration column with 50 mM Tris pH 7.5, 150 mM NaCl and 1 mM DTT. Purified proteins were flash frozen and stored at -80 °C.

#### S4. Cell culture and transfections

HEK 293T cells were obtained from the Crick Cell Services and cultured in DMEM (Dulbecco's Modified Eagle's Medium, Gibco, D6429) supplemented with 10% (v/v) fetal bovine serum (Gibco, A52567) and penicillin-streptomycin antibiotics (Gibco, 11548876). Prior to transfection, cells were seeded into 10 cm dishes for 24 hours. Transfections were performed with Lipofectamine 2000 (Invitrogen, 11668019) diluted with Opti-MEM (Gibco, 31985070), following the manufacturer's instructions. Per each 10 cm dish, 3 µg (LgBiT-Flag) or 12 µg (BRD3-HiBiT/BRD3-V5) DNA was transfected. These quantities were scaled for transfections with smaller dish sizes. Cells were washed with PBS and harvested 24 h post-transfection by scraping into PBS. Cells were pelleted at 800 *x g* for 2 min in a cooled benchtop centrifuge, and cell pellets were either lysed immediately for use in downstream assays or flash frozen and stored at -80 °C.

#### S5. Protein quantitation with NanoBiT

*HiBiT quantitation:* A standard curve for HiBiT quantitation was prepared by incubating LgBiT protein (1:200, N401A, Promega), furimazine substrate (1:100, N246A, Promega) and a dilution series of HiBiT-HaloTag control protein (Promega, N3010) in PBS-T (PBS, 0.1 % Tween20) in a white 96-well plate (M0187, Greiner) with a final well volume of 100 µL. The plate was incubated at RT for 10 min with gentle rocking and read using a CLARIOstar PLUS Plate Reader with a 470 nm (80 nm bandpass) filter.

*LgBit quantitation:* A standard curve for LgBiT quantitation was prepared by incubating HiBiT control protein (100 nM, N3010, Promega), furimazine substrate (1:100, N246A, Promega) and a dilution series of LgBiT protein (N401A, Promega) in PBS-T in a white 96-well plate (M0187, Greiner) with a final well volume of 100 µL. The plate was incubated and read as for HiBiT quantitation above.

#### S6. Immunoprecipitation trials

*Bead preparations:* V5 and HiBiT antibody-coated beads were prepared by mixing Protein G Dynabeads (50 µL) with 2 µg anti-V5 (Cell Signalling technology, 13292S) or anti-HiBiT (Promega, CS2006A01) in PBS-T at RT for 10 min with rotation. Magnetic beads for Flag immunoprecipitations were purchased from Sigma (M8823).

*Cell lysis:* Cell pellets were lysed with cold NP40 lysis buffer (25 mM Tris-HCl, pH 7.5, 150 mM NaCl, 5 mM $MgCl_2$, 1% NP-40, 5% glycerol) supplemented with cOmplete™ EDTA-free Protease Inhibitor

Cocktail (Roche) (1 mL lysis buffer per 10 cm dish confluent cells). Samples were vortexed and incubated on ice for 10 min. Lysates were clarified by centrifugation at 17,000 *x g* for 15 min in a cooled benchtop centrifuge.

*Immunoprecipitations:* Lysates were incubated with varying quantities of magnetic beads prepared as above and incubated at 4 °C for 30 min with rotation. Supernatants post-incubation were retained for gel analysis, and beads were washed 3x with PBS-T. Bead (IP) samples and supernatants were heated to 95 °C for 5 min in 1x laemmli sample buffer and analysed by western blotting (see section S7). Band intensities were quantified using Fiji's gel analysis tool.[2] Additionally, the amounts of immunoprecipitated LgBiT and BRD3-HiBiT proteins were quantified by luminescence assays as described in S5.

#### S7. Immunoblotting

Samples in 1x laemmli buffer (BioRad, 1610747) containing 1.4 M beta-mercaptoethanol were resolved by electrophoresis on a 4-20% Bis-Tris Precast gel (Millipore mPAGE, MP42) in 1x MOPS SDS running buffer (Invitrogen, NP0001). Proteins were transferred to a PVDF membrane (Amersham Hybond 0.45, 10600023) by wet transfer with a BioRad Mini Trans-Blot Cell. Membranes were blocked with 5% milk in PBS with 0.1% Tween20 for 1 h at room temperature, followed by incubation with the following primary antibodies at the indicated dilutions; mouse anti-HiBiT (N7200, Promega, 1:2500), rabbit anti-BRD3 (A302-367A, Bethyl Laboratories, 1:2500), rabbit anti-V5 (#13202, Cell Signalling Technologies, 1:000), rabbit anti-BRD2 (A302-583A, Bethyl Laboratories, 1:2500), rabbit anti-BRD4 (HPA061646, Sigma, 1:2500), anti-Flag HRP (A8592, Sigma, 1:10000).

#### S8. mRNA synthesis

##### S8.1 Individual templates

mRNA templates M_HiBiT and BRD3.1b[3] were constructed by two rounds of overlapping PCR and purified as described previously,[4] using the template primers detailed in Table S1.

##### S8.2 Library synthesis

NNK6-12 libraries for BRD3 selections were prepared as described previously.[4] The 11mer randomised library for LgBiT selections was prepared as follows in a two-step PCR. The first PCR was performed on a 100 µL scale (1x KOD polymerase buffer, 1 mM $MgCl_2$, 0.2 mM dNTPs, 0.6 µM T7g10M.F46 primer, 0.5 µM 1aa_linear primer, 0.8 µL KOD polymerase), with an annealing temperature of 55 °C for 7 cycles. The second PCR was performed on a 1 mL scale using the product from the first round (100 µL) as a template (1x KOD polymerase buffer, 1 mM $MgCl_2$, 0.2 mM dNTPs, 0.25 µM T7g10M.F46 primer,

0.25 µM GS3an13.R36 primer, 8 µL KOD polymerase), with an annealing temperature of 61 °C for 20 rounds.

The PCR product was purified by phenol-chloroform extraction followed by ethanol precipitation. The purified product was then transcribed overnight using T7 RNA polymerase (Thermo Scientific) following the manufacturer's protocol. The RNA was isolated by isopropanol precipitation and further purified by urea denaturing 8% PAGE gel (19:1 acrylamide/bis-acrylamide).

#### S9. Quantifying RNA/DNA stability in HEK 293T cell lysates

An NNK9 mRNA library was reverse transcribed with M-MLV RTase, RNase H minus as described previously.[5] The mRNA/cDNA was incubated with HEK 293T lysate at 4 °C, prepared with NP40 lysis buffer supplemented with cOmplete™ EDTA-free Protease Inhibitor Cocktail (Roche) (1 mL lysis buffer per 10 cm dish confluent cells). At various time points, samples from the mix were quenched by boiling at 95 °C for 5 min in PCR buffer. DNA recovery as a percentage of the input DNA was assessed by quantitative real-time PCR using primers T7g10M.F46 and CGS3an13.R22 (Table S4).

#### S10. Aminoacylation of tRNAs

Flexizymes (eFx and dFx), initiator $tRNA^{fMet}_{CAU}$ and elongator $tRNA^{Asn}_{CAU}$ were synthesised according to the previously published protocol.[6]

*CME substrates:* Aminoacylation was performed by mixing 5 mM ClAc-D-Tyr-CME or ClAc-L-Trp-CME with 600 mM $MgCl_2$, 20% DMSO, 25 µM eFx and 25 µM initiator $tRNA^{fMet}_{CAU}$ and 50 mM HEPES-KOH (pH 7.5), following a previously published protocol.[6] The mixture was incubated for 2 h on ice. The resulting aminoacyl-microhelix/tRNA was purified by ethanol precipitation. Pellets were washed twice with 70% ethanol containing 0.1 M sodium acetate (pH 5.2), frozen with liquid $N_2$ and stored at -80 °C.

*DBE substrates:* aminoacylations were performed as for CME substrates with the following reaction mixture: 5 mM *N*-methyl-alanine-DBE or acetyllysine-DBE,[3] 600 mM $MgCl_2$, 20% DMSO, 25 µM dFx and 25 µM elongator $tRNA^{Asn}_{CAU}$ and 50 mM HEPES-KOH (pH 7.5).

#### S11. Mock selection with purified and cell lysate BRD3

Translation of control peptide 3.1b (MKTIMGMTWRTMQCGSGSGS) was performed using a PURExpress$^{TM}$ (ΔRF123) *in vitro* protein synthesis kit (NEB, E6850S) as described previously.[4] Briefly, translation mixtures were prepared on ice by combining 0.78 µL solution A (50 mM HEPES-KOH, 2 mM ATP (Jena Bioscience), 2 mM GTP (Jena Bioscience), 1 mM CTP (Jena Bioscience), 1 mM UTP (Jena Bioscience), 20 mM creatine phosphate, 100 mM potassium acetate, 2 mM spermidine, 6 mM magnesium acetate, 1.5 mg/ml *E. Coli* tRNA (Roche), 14 µM DTT), 1.5 µL solution B (PURExpress$^{TM}$), 0.5 µL each aminoacyl-tRNA (250 µM, ClAc-Trp-$tRNA^{fMet}_{CAU}$ and AcK-$tRNA^{Asn}_{CAU}$, prepared as described above S10.), 0.5 µL mRNA template (BRD3.1b, 10 µM), 0.5 µL amino acid mixture (19 aa -Met, 5 mM each), 0.72 µL water. Methionine and 10-formyl-5,6,7,8-tetrahydrofolic acid were not included in the translations to enable reprogramming of Met codons. The translation reaction mixture was incubated at 37 °C for 1 h. Following addition of 200 mM EDTA (pH 8.0), the translated mixture was reverse transcribed with M-MLV RTase, RNase H minus.

Mock selection with recombinant BRD3-BD1: An equivalent volume of 2x blocking buffer was added to the translation mixture (50 mM HEPES pH 7.5, 150 mM NaCl, 2 mM DTT, 0.1% Tween20, 0.2% (w/v) acetylated bovine serum albumin). M280 streptavidin Dynabeads (Invitrogen, 11205D) with immobilised BRD3-BD1 (200 nM, positive selection) or with biotin (negative selection) were incubated with translation mixtures (10 µL) on ice for 30 min.

Mock selection with BRD3-HiBiT from HEK 293T cell lysates: HEK 293T cells transfected with BRD3-HiBiT were lysed with NP40 lysis buffer, as described in section S4 and S6. HiBiT antibody-coated beads were prepared by incubating Protein G Dynabeads (Invitrogen, 10003D) with anti-HiBiT antibody (Promega, 30E5) in PBS-T at RT for 10 min with rotation. HiBiT antibody-coated beads were incubated with BRD3-HiBiT lysate (positive selection) or NP40 lysis buffer (negative selection) at 4 °C for 30 min with rotation. Translation mixtures were added to positive beads (~140 nM BRD3-HiBiT, estimated by HiBiT quantitation assay – see section S5) and negative beads, and incubated at 4 °C for 30 min with rotation.

Both mock selections: Positive and negative selection beads were each washed with 3 x 20 µL with ice-cold assay buffer (50 mM HEPES pH 7.5, 150 mM NaCl, 2 mM DTT, 0.1% Tween20). PCR buffer (100 µL) was added to the beads and the retained peptide-mRNA/DNA hybrids were eluted by heating to 95 °C for 5 min. DNA recovery as a percentage of the input DNA was assessed by quantitative real-time PCR using primers T7g10M.F46 and CGS3an13.R22 (Table S4).

#### S12. RaPID selections

RaPID screens were adapted from protocols previously described.[5]

##### 12.1 LgBiT selections

An 11mer randomised mRNA library was prepared as described in section S8.2. Approximately 100 copies of the M_HiBiT mRNA template were spiked into the library. The puromycin-ligated library (10 pmol) was *in vitro* translated on a 10 µL scale (30 min, 37 °C then 12 min, 25 °C) using solution A (section S11) supplemented with 100 µM 10-formyl-5,6,7,8-tetrahydrofolic acid and an amino acid mix containing all 20 amino acids, as no reprogramming was used. Following addition of 200 mM EDTA (pH 8.0) and incubation at 37 °C for 30 min, the translated mixture was reverse transcribed with M-

MLV RTase, RNase H minus. The resulting mixture was split into two for the two selection conditions described below.

*Purified LgBiT selection:* Translation mix (10 µL) was mixed with 2x blocking solution (10 µL, 50 mM HEPES pH 7.5, 150 mM NaCl, 0.1 % Tween20, 2 mM DTT, 0.2 % w/v acetyl-BSA). The library was added to M280 Streptavidin Dynabeads (Invitrogen, 11205D) with immobilised biotinylated LgBiT (200 nM) and incubated at 4 °C with rotation for 1 h. Beads were washed with 3 x 20 µL 50 mM HEPES pH 7.5, 150 mM NaCl, 0.1 % Tween20, 2 mM DTT and resuspended in 100 µL PCR buffer.

*LgBiT cell lysate selection:* Translation mix (10 µL) was mixed with 2x blocking solution (10 µL, HEK 293T lysate, 0.2 % w/v acetyl-BSA) and HEK 293T lysate transfected with LgBiT at ~200 nM. The library was added to Anti-Flag M2 magnetic beads (3 µL, Sigma, M8823) and incubated at 4 °C with rotation for 1 h. Beads were washed with 3 x 20 µL NP40 lysis buffer and resuspended in 100 µL PCR buffer.

*Both selections:* Bound peptide-mRNA/cDNA hybrids were eluted from the beads by boiling at 95 °C for 5 min. Library enrichment was assessed by quantitative real-time PCR relative to the input DNA library using primers T7g10M.F46 and CGS3an13.R22 (Table S4). Recovered DNA was amplified by PCR and used as the input for the next selection round. This cDNA was transcribed into mRNA using T7 RNA polymerase (Thermo Scientific, 18033019), following the manufacturer's instructions. A total of five rounds of selection were performed for each condition, as above, with translations performed on a 5 µL scale. From Round 2 onwards, negative selections were also included. The libraries in blocking buffer were incubated with 3 x 3 µL Flag (cell lysate) or streptavidin (purified protein selection) beads for 30 min at 4 °C with rotation, to deplete the libraries of sequences binding to the beads or antibody. Following the negative selections, the positive selections were performed as above, with HEK 293T lysate transfected with 3xFlag LgBiT (~200 nM) being added to the mixture at this stage for the cell lysate selection.

Following five rounds of RaPID selection, double indexed libraries (Nextera XT indices) were prepared from library DNA recovered from positive and negative selections from all rounds and sequenced on the Illumina HiSeq 4000 or NovaSeq platform with single-ended 100 bp reads. Each DNA sequence was converted to a peptide sequence and ranked by total read number - see Supplementary Files 1 & 2.

#### 12.2 BRD3 selections

*Both selections, Round 1:* An NNK6-12 mRNA library was prepared as described previously.[4] The puromycin-ligated mRNA library (10 µL, 50 pmol) was *in vitro* translated on a 50 µL scale (30 min, 37 °C then 12 min, 25 °C) using 7.8 µL solution A (section S11), 15 µL solution B (PURExpress$^{TM}$, ΔRF123),

5 µL 19 amino acid mix (-Met, 5 mM each) and 2.5 µL each aminoacyl-tRNA (250 µM, ClAc-D-Tyr-$tRNA^{fMet}_{CAU}$ and *N*-methyl-Ala-$tRNA^{Asn}_{CAU}$, prepared as described in S10.) and 7.2 µL water. Methionine and 10-formyl-5,6,7,8-tetrahydrofolic acid were not included in the translations to enable reprogramming of Met codons. Following addition of 200 mM EDTA (pH 8.0) and incubation at 37 °C for 30 min, the translated mixture was reverse transcribed with M-MLV RTase, RNase H minus.

*BRD3-HiBiT selection 1, Round 1:* The reverse transcribed library was mixed with blocking buffer (HEK 293T lysate + 0.2% w/v acetyl-BSA) to give a final volume of 100 µL. A HEK 293T cell pellet transfected with BRD3-HiBiT (see section S4) was lysed with NP40 lysis buffer supplemented with cOmplete™ EDTA-free Protease Inhibitor Cocktail (1 mL). BRD3-HiBiT was immunoprecipitated with 80 µL HiBiT antibody-coated Protein G Dynabeads (prepared as in S6), which was estimated to contain a BRD3-HiBiT concentration of 200 nM following resuspension in 100 µL RaPID library. The library was incubated with BRD3-HiBiT immunoprecipitation beads at 4 °C for 30 min with rotation. The beads were washed with 3 x 200 µL NP40 lysis buffer, changing tubes each time. Retained peptide-mRNA/cDNA complexes were eluted from the beads by boiling at 95 °C for 5 min. DNA recovery as a percentage of the input DNA was assessed by quantitative real-time PCR using primers T7g10M.F46 and CGS3an13.R22 (Table S4). The recovered DNA was amplified by PCR and used as the input for the next round of selection.

*BRD3-HiBiT selection 1, Rounds 2-5:* mRNA was transcribed from the cDNA recovered from the previous round using T7 RNA polymerase (Thermo Scientific, 18033019), following the manufacturer's instructions. The library was ligated to puromycin, then a translation was performed, as for Round 1, on a 5 µL scale with 5 pmol puromycin-ligated mRNA. Following addition of 200 mM EDTA (pH 8.0) and incubation at 37 °C for 30 min, the translated mixture was reverse transcribed with M-MLV RTase, RNase H minus. The library was mixed (1:1) with blocking buffer (HEK 293T lysate + 0.2% w/v acetyl-BSA) to give a final volume of 18.88 µL.

The peptide-mRNA/cDNA library was incubated with 5 x 4 µL HiBiT antibody-coated beads for 10 min at 4 °C with rotation, to remove sequences binding to the immobilised antibody in negative selections. A HEK 293T cell pellet transfected with BRD3-HiBiT (see section S4) was lysed with NP40 lysis buffer supplemented with cOmplete™ EDTA-free Protease Inhibitor Cocktail (50 µL) and immunoprecipitated with 4 µL HiBiT antibody-coated beads for 30 min at 4 °C with rotation (see section S6). The library was added to the immunoprecipitation beads and incubated for 30 min at 4 °C with rotation. Positive and negative selection beads were washed with 3 x 20 µL cold NP40 lysis buffer and resuspended with 100 µL PCR mix. Retained peptide-mRNA/cDNA complexes were eluted from the beads by boiling at 95 °C for 5 min. DNA recovery as a percentage of the input DNA was assessed by quantitative real-time PCR using primers T7g10M.F46 and CGS3an13.R22 (Table S4). The recovered DNA from the positive selection was amplified by PCR and used as the input for the next round of selection. The process was repeated until a total of five selection rounds had been performed.

Following five rounds of RaPID selection, double indexed libraries (Nextera XT indices) were prepared from library DNA recovered from positive and negative selections from all rounds and sequenced on the Illumina HiSeq 4000 or NovaSeq platform with single-ended 100 bp reads. Each DNA sequence was converted to a peptide sequence and ranked by total read number - see Supplementary File 3.

*BRD3-HiBiT and BRD3-V5 selection 2, Round 1:* The reverse-transcribed library was mixed with HEK 293T cell lysate (non-transfected), 2 mg/mL salmon sperm DNA and 15 µL 2% w/v acetyl-BSA to a final volume of 150 µL. Extensive negative selections were performed with a 1:1:1 mixture of HiBiT and V5 antibody-coated Protein G Dynabeads and Flag M2 magnetic beads (5 x 48 µL, 30 min at 4 °C with rotation). BRD3-HiBiT HEK 293T cell pellets were lysed with NP40 lysis buffer supplemented with cOmplete™ EDTA-free Protease Inhibitor Cocktail (1 mL). Lysate was incubated with 80 µL HiBiT antibody-coated beads at 4 °C with rotation for 30 mins (see section S6). The beads were washed with 3 x 200 µL NP40 lysis buffer, changing tubes each time. Retained peptide-mRNA/cDNA complexes were eluted from the beads by boiling at 95 °C for 5 min. DNA recovery as a percentage of the input DNA was assessed by quantitative real-time PCR using primers T7g10M.F46 and CGS3an13.R22 (Table S4). The recovered DNA was amplified by PCR and used as the input for the next round of selection.

*BRD3-HiBiT selection 2, Rounds 2-5:* The next four rounds of selection were performed as in BRD3-HiBiT selection 1, with the following alterations. 2 mg/mL salmon sperm DNA was included in the blocking buffer, 200 µL transfected lysate was used for immunoprecipitation, greater quantities of beads were used for immunoprecipitation (8 µL HiBiT and 10 µL V5 antibody-coated beads), and negative selections were performed with a mixture of HiBiT, V5 and Flag magnetic beads (3 x 9 µL mixed beads, 30 min at 4°C with rotation). BRD3-HiBiT transfected pellets and HiBiT immunoprecipitations were used for positive selections in Rounds 3 & 5, while BRD3-V5 transfected pellets and V5 immunoprecipitations were used in Rounds 2 & 4.

Following five rounds of RaPID selection, double indexed libraries (Nextera XT indices) were prepared from library DNA recovered from positive and negative selections from all rounds and sequenced on the Illumina HiSeq 4000 or NovaSeq platform with single-ended 100 bp reads. Each DNA sequence was converted to a peptide sequence and ranked by total read number - see Supplementary File 4.

#### 12.3 Analysis of selection data

Sequence alignments were produced using CLC Main Workbench (Qiagen). Logo plots were produced using Berkley's WebLogo server.[7]

### S13. Peptide synthesis

Peptides with an amidated C-terminus were synthesised by standard Fmoc-strategy solid-phase chemistry using an automated peptide synthesizer (Gyros Prelude X) with Rink Amide MBHA resin LL (Novabiochem, 0.05 mmol scale). Couplings were performed with Oxyma pure (5 eq), diisopropylcarbodiimide (5 eq) and Fmoc-protected amino acids (5 eq) in dimethylformamide (DMF). Couplings were performed at 25 °C for His, 60 °C for Cys and 90 °C for all other amino acids. Following the final coupling, the N-terminal Fmoc group was removed and the N-terminal amine was capped by acetylation (10% acetic anhydride in DMF, 1 h, RT) or chloroacetylation (5 eq *N*-(chloroacetoxy)succinimide in DMF, 1 h, RT) for thioether-cyclised peptides.

Peptides were cleaved from the resin with trifluoroacetic acid (TFA, 92.5%), $H_2O$ (2.5%), 2,2'-(Ethylenedioxy)diethanethiol (2.5 %) and triisopropylsilane (2.5%) at RT for 2-4 hours. Peptides were precipitated with cold diethyl ether, washed 3 times and dried. Crude peptides were dissolved in DMSO, and cyclised by addition of triethylamine to pH >8.0 (2 h, RT) if chloroacetylated.

Linear and cyclic peptides were purified by reverse-phase HPLC (Waters XBridge prep BEH C18, 19 x 250 mm, 10 µm) and analysed by analytical HPLC (Waters XBridge BEH C18, 4.6 x 250 mm, 5 µm), both systems running gradients of water (0.1% v/v TFA) and acetonitrile (0.1% v/v TFA). Peptides were additonally analysed by LCMS (Aquity UPLC BEH C18, 2.1 x 50 mm, 1.7 µm). Analytical HPLC traces for all peptides can be found in Figures S13-14 and LC/MS analysis is in Table S1-2. Purified peptides were reconstituted in DMSO and their concentrations estimated using calculated extinction coefficients at 280 nm.

### S14. Click reactions

TAMRA-labelled peptides: Purified, azide-containing peptides (1 µmol, ~ 2 mg) were dissolved in DMSO and added to TAMRA-alkyne (1.5 eq, prepared at 10 mg/mL in DMSO, Sigma, 900932), sodium ascorbate (2.5 eq, prepared at 25 mM in $H_2O$) and $CuH_2SO_4$: Tris(3-hydroxypropyltriazolylmethyl)amine (1:1 in $H_2O$ at 10 mM, 1 eq) in a foil-wrapped glass vial.

Biotinylated peptides: Biotinylated peptides were prepared using the same click conditions as TAMRA-labelled peptides above, substituting biotin-$PEG_4$-alkyne (1.5 eq, prepared at 10 mg/mL in DMSO, Sigma, 1262681-31-1) for TAMRA-alkyne.

Click reactions were incubated at RT for 2 h, purified by preparative HPLC and analysed by analytical HPLC and LC/MS as above. See Figure S15 for analytical HPLC traces and Table S3 for masses observed by LC/MS.

#### S15. Luminescence complementation assays

##### S15.1 LgBiT peptide direct binding assays

Direct binding of HiBiT-HaloTag control protein (Promega, N3010) and M_HiBiT peptide were measured by mixing a dilution series of the HiBiT-protein/peptide with LgBiT protein (100 pM, N401A, Promega) and furimazine substrate (1:100, N246A, Promega) in PBS-T in a white 96-well plate (M0187, Greiner) with a final well volume of 100 µL. The plate was incubated at RT for 10 min with gentle rocking and read using a CLARIOstar PLUS Plate Reader with a 470 nm (80 nm bandpass) filter.

##### S15.2 HiBiT-HaloTag competition assays

Each well of a 96-well white plate (M0187, Greiner) was prepared with HiBiT-HaloTag control protein (24 nM, Promega, N3010), LgBiT protein (100 pM, N401A, Promega) and furimazine substrate (1:100, N246A, Promega). Dilution series of LgBiT peptides for competition were prepared in PBS-T and added to the plate for a final well volume of 100 µL. Plates were incubated and measured as for direct binding assays.

#### S16. NanoBRET assays

##### S16.1 Direct binding with fluorescent peptides

Cell pellets from HEK 293T cells transfected with HiBiT-tagged constructs were suspended with NP40 lysis buffer (1 mL per 10 cm dish – 25 mM Tris-HCl, pH 7.5, 150 mM NaCl, 5 mM $MgCl_2$, 1% NP-40, 5% glycerol) supplemented with cOmplete™ EDTA-free Protease Inhibitor Cocktail (Roche), vortexed and incubated on ice for 10 min. Lysates were clarified by centrifugation at 17,000 *x g*, 4 °C for 15 min. Lysates were diluted with NP40 lysis buffer and mixed with LgBiT protein (1:200, N401A, Promega), furimazine substrate (1:100, N246A, Promega) in white 96-well plates (M0187, Greiner). Dilution series of TAMRA-labelled peptides were prepared in PBS-T and added to the plate for a final well volume of 100 µL. Plates were incubated on a rocker at RT for 30 min before reading with a CLARIOstar PLUS Plate Reader with 460 nm (80 nm bandpass) and 580 nm (30 nm bandpass) filters, with gains of 3600 for each. BRET ratios were calculated by dividing the fluorescence intensity by the luminescence intensity.

##### S16.2 NanoBRET competition assays

Lysates were prepared as above. Each well of a 96-well white plate (M0187, Greiner) was prepared with diluted HiBiT-containing lysate, LgBiT protein (1:200, N401A, Promega), furimazine substrate

(1:100, N246A, Promega) and a TAMRA-labelled peptide for competition (concentration indicated in figure legends). Dilution series of non-labelled peptides for competition were prepared in PBS-T and added to the plate for a final well volume of 100 µL. Plates were incubated and measured as for direct binding assays.

#### S17. Surface plasmon resonance (SPR)

SPR experiments were carried out using a Biacore S200 and a Biotin CAPture kit, series S (Cytiva). Experiments were performed at 25 °C in 10 mM HEPES pH 7.4, 150 mM NaCl, 0.005% surfactant P20, 1 mM DTT and 0.1 % DMSO. His-bio-BRD3-ET (100 nM) and His-bio-BRD3-L (1000 nM) were immobilised to RU ~350 and ~1000 respectively at a flow rate of 10 µL/s. Five-point trebling dilutions series were prepared for each peptide, and single-cycle kinetic experiments were performed with a 120 s contact time and 360 s dissociation time. Between cycles, the chip was regenerated following the manufacturer's protocols. Kinetic and affinity data were analysed using Biacore S200 Evaluation software (Cytiva). Data were fitted to a 1:1 binding model.

#### S18. Preparation of nuclear extracts

Cell pellets were resuspended in cold cytoplasmic lysis buffer (10 mM HEPES pH 8.0, 1.5 mM $MgCl_2$, 10 mM KCl, 1 mM EDTA, 0.05% NP40) supplemented with cOmplete™ EDTA-free protease inhibitor cocktail, vortexed briefly and incubated on ice for 30 min. Nuclear material was pelleted at 800 *x g* for 2 min, the supernatant discarded, pellets resuspended with cold nuclear extraction buffer (5 mM HEPES pH 8.0, 1.5 mM $MgCl_2$, 300 mM, 0.2 mM EDTA, 25% glycerol and protease inhibitors) and incubated on ice for 30 min with regular vortexing. Lysates were clarified by centrifugation (15 min, 13,000 *x g*, 4 °C) and diluted with nuclear extraction buffer lacking NaCl to reduce salt concentration to 150 mM.

#### S19. Pulldowns with biotinylated peptides

Dynabeads™ M280 Streptavidin (11205D, Invitrogen) for pulldowns were prepared by incubating beads with an excess of biotin or biotinylated peptides in PBS-T at RT for 15 min, and washed with PBS-T. Pulldowns were performed using a KingFisher Duo Prime Purification system (ThermoScientific). Briefly, magnetic beads coated with biotin or biotinylated peptides were incubated with HEK 293T nuclear extracts for 1 h with mixing, followed by three brief PBS-T washes and elution with 1x laemmli sample buffer containing 1.4 M beta-mercaptoethanol. Samples were analysed by western blotting.

### Supplementary Figures

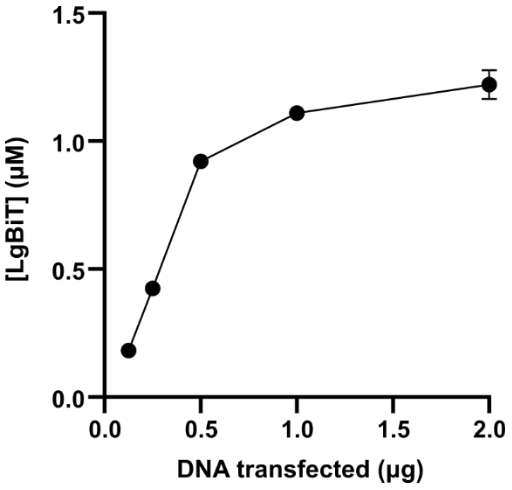

**Figure S1:** LgBiT protein quantified from cell lysates using HiBiT complementation luminescence assays following transfection of varying quantities of 3xFlag-LgBiT into HEK 293T cells. Transfections were performed 16 h after seeding 9 x $10^5$ HEK 293T cells into wells of a 6-well plate. Dishes were lysed 24 h post-transfection with 160 µL lysis buffer prior to quantitation.

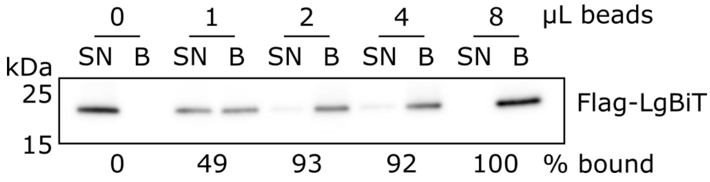

**Figure S2.** Flag immunoprecipitation of LgBiT from HEK 293T cell lysates. Bands were quantified using the gel analysis tool in Fiji. Bead quantities shown were used to immunoprecipitate 200 nM LgBiT from 40 µL lysate. Supernatant (SN) and bead (B) samples were normalised to the same volume before western blot to allow quantitation of the bead-bound fraction, % bound = B / (SN+B).

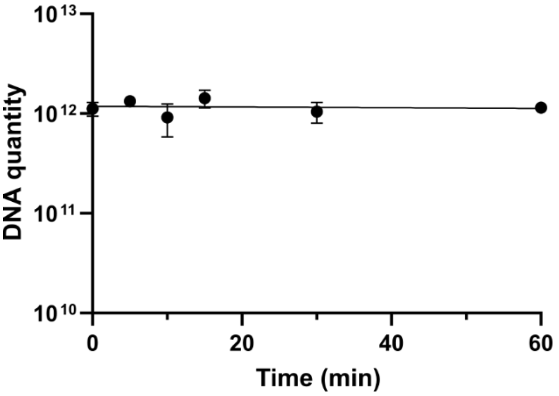

**Figure S3.** mRNA/cDNA stability in a HEK 293T cell lysate. A reverse-transcribed NNK12 library (5 µL, 2 µM) was incubated with 15 µL HEK 293T lysate (prepared by lysing a confluent 10 cm dish with 1 mL lysis buffer containing protease inhibitors). Samples were removed at various timepoints and boiled at 95 °C for 5 min to denature lysate proteins prior to qPCR quantitation.

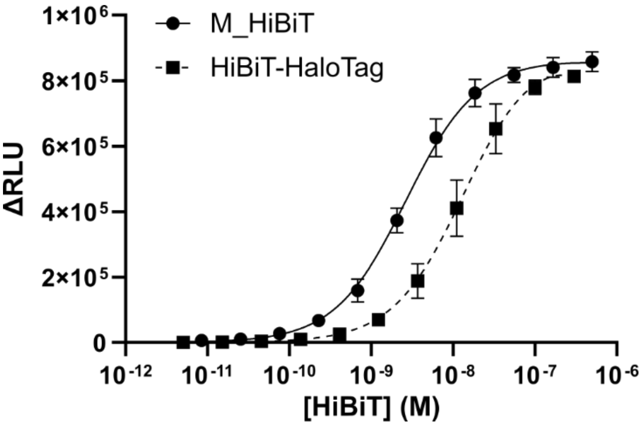

**Figure S4.** Luminescence complementation assays to measure the direct binding of M_HiBiT peptide and HiBiT-HaloTag control protein to recombinant LgBiT protein. $K_D$ M_HiBiT = 2.7 ± 0.2 nM and $K_D$ HiBiT-HaloTag = 14 ± 2 nM (n = 4). RLU – relative luminescence units.

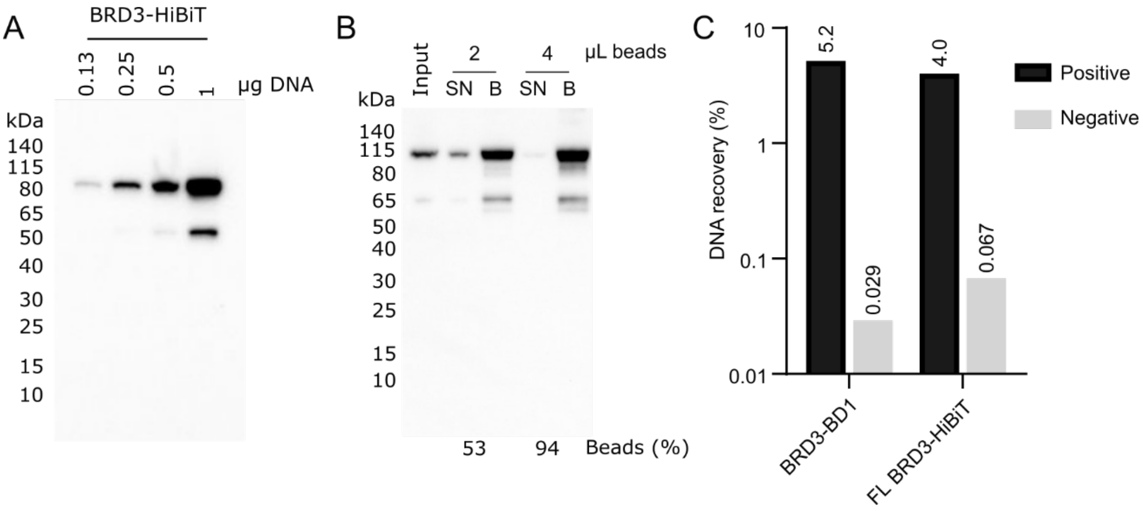

**Figure S5. HiBiT-BRD3 expression trials and test immunoprecipitations. A.** Expression trials with BRD3-HiBiT. Western blot probed with anti-HiBiT antibody showed band at the expected size (81 kDa). Transfections were performed 16 h after seeding 8 x $10^5$ HEK 293T cells into wells of a 6-well plate. Dishes were lysed 24 h post-transfection with 160 µL lysis buffer prior to quantitation. **B.** Immunoprecipitation trial to quantify volume of HiBiT antibody-coated beads required to capture 10 nM BRD3-HiBiT from 100 µL HEK 293T lysate (quantified using HiBiT luminescence assay). SN – supernatant, B – beads. Supernatant (SN) and bead (B) samples were normalised to same volume before western blot to allow quantitation of the bead-bound fraction, % bound = B / (SN+B). Band intensities were quantified using Fiji's gel analysis tool. **C.** DNA recovery of a BRD3-BD1 cyclic peptide (BRD3.1b - cyclic-WKTIKGKTWRTKQC(S-)GSGSGS,[8] K = acetyl-K) in a mock selection using biotinylated BRD3-BD1 or full length BRD3-HiBiT immunoprecipitated from HEK 293T cell lysates.

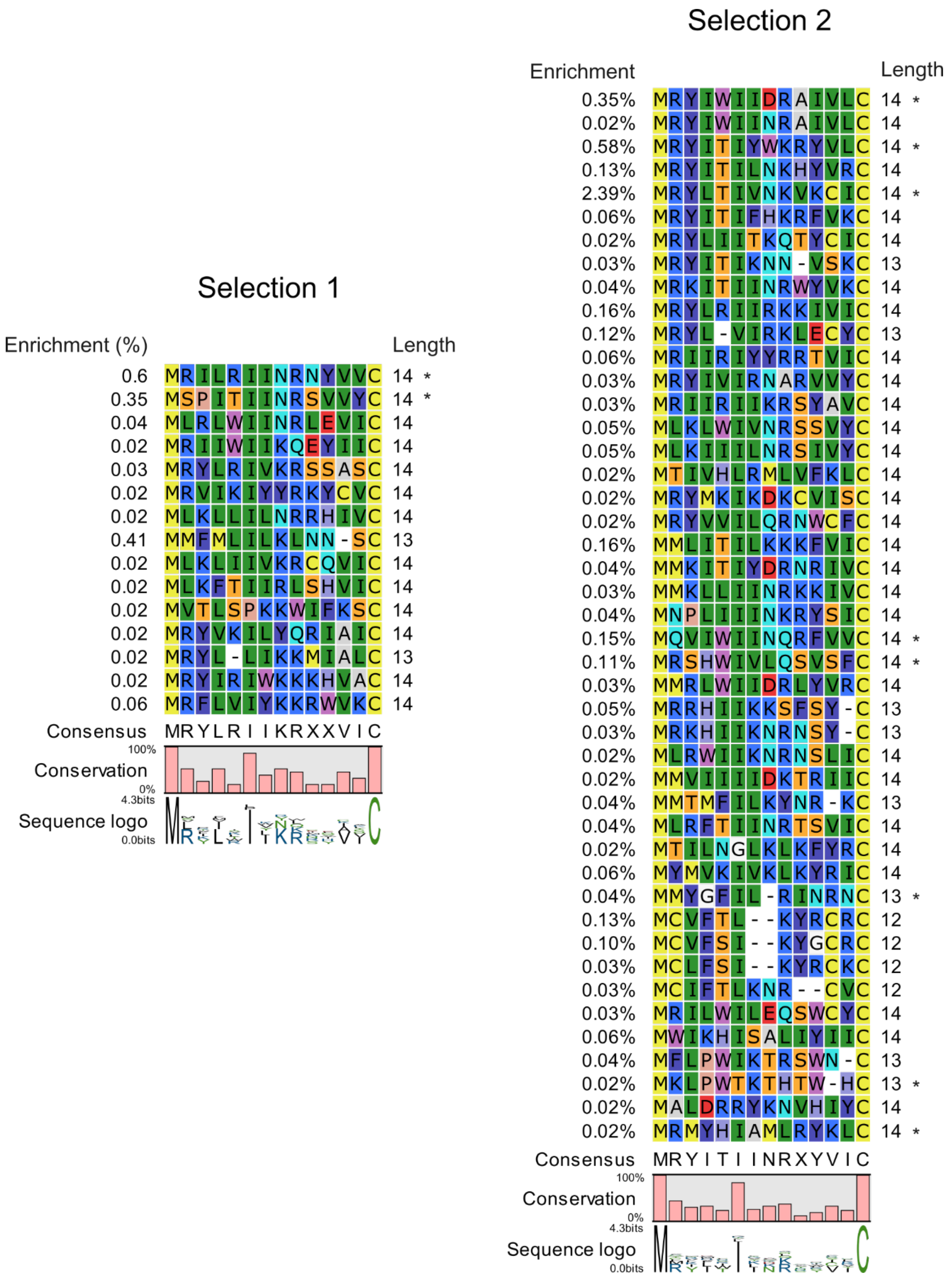

**Figure S6. BRD3 positive hits.** Alignment of sequences only enriched in the positive selections against BRD3 (not enriched in negative selections). Shown are sequences appearing with a frequency of >0.01%. Sequences marked with * were selected for synthesis.

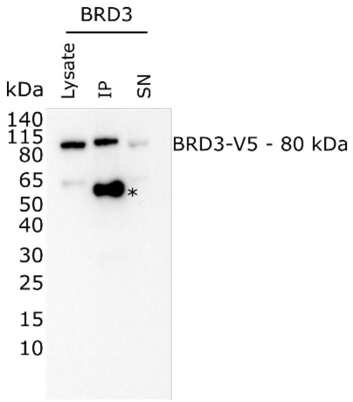

**Figure S7.** BRD3-V5 expression and immunoprecipitation trial. BRD3-V5 was immunoprecipitated from 180 µL transfected lysate with 10 µL V5 antibody-coated beads. IP – immunoprecipitation, SN – supernatant. The band denoted * is believed to be the antibody heavy chain detected by the secondary antibody.

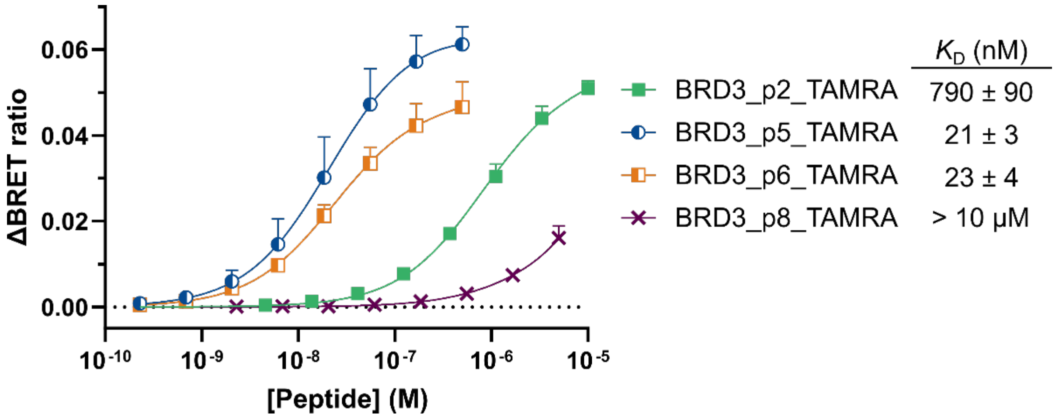

**Figure S8.** NanoBRET direct binding assays with full-length BRD3-HiBiT and TAMRA-labelled peptides. The data displayed are the mean values + one standard error of the mean from at least three independent replicates.

#### BRD3-ET

Kinetics | Affinity

BRD3_p1

BRD3_p2

BRD3_p3.1

BRD3_p3.2

BRD3_p4

BRD3_p5

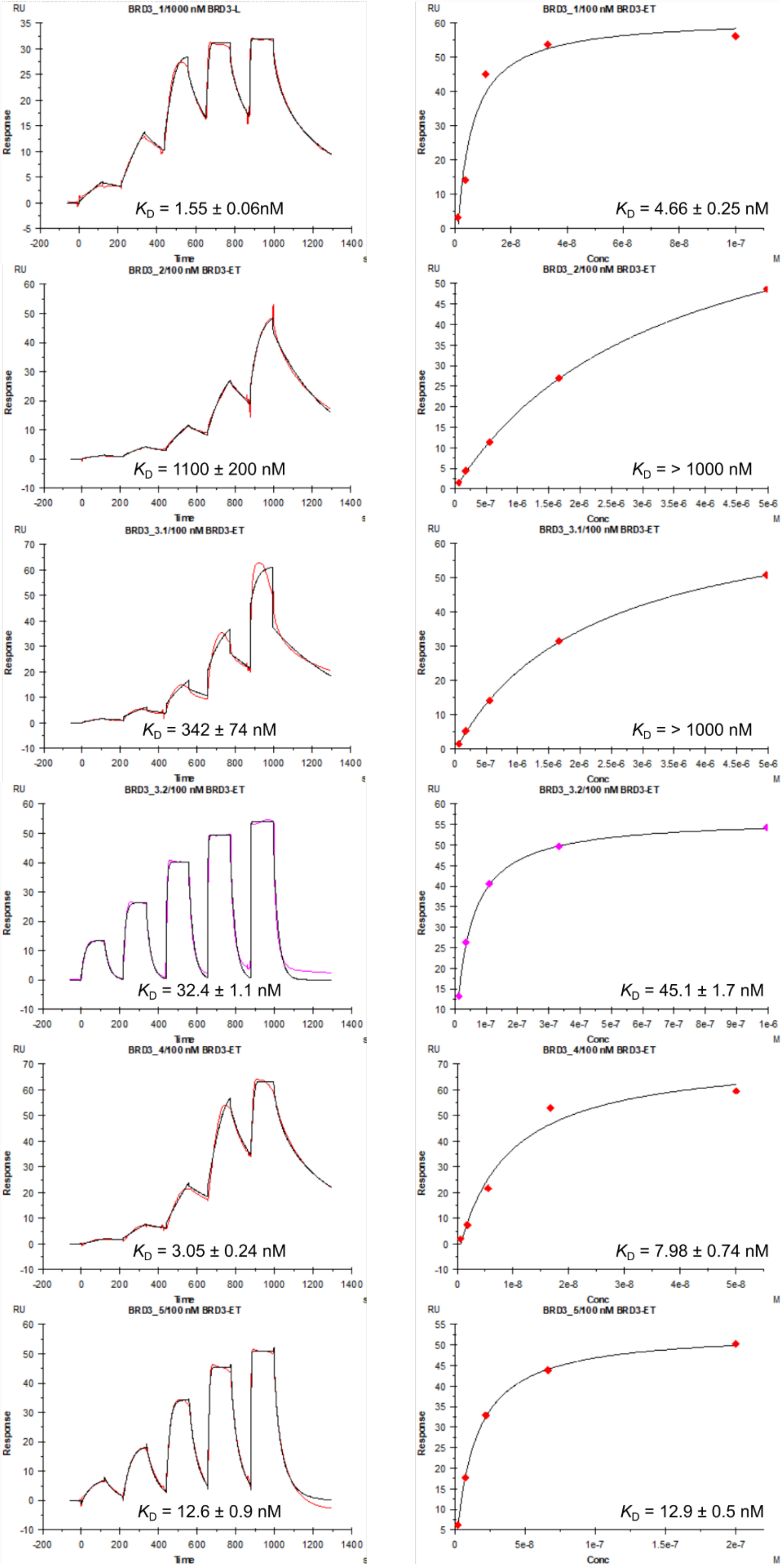

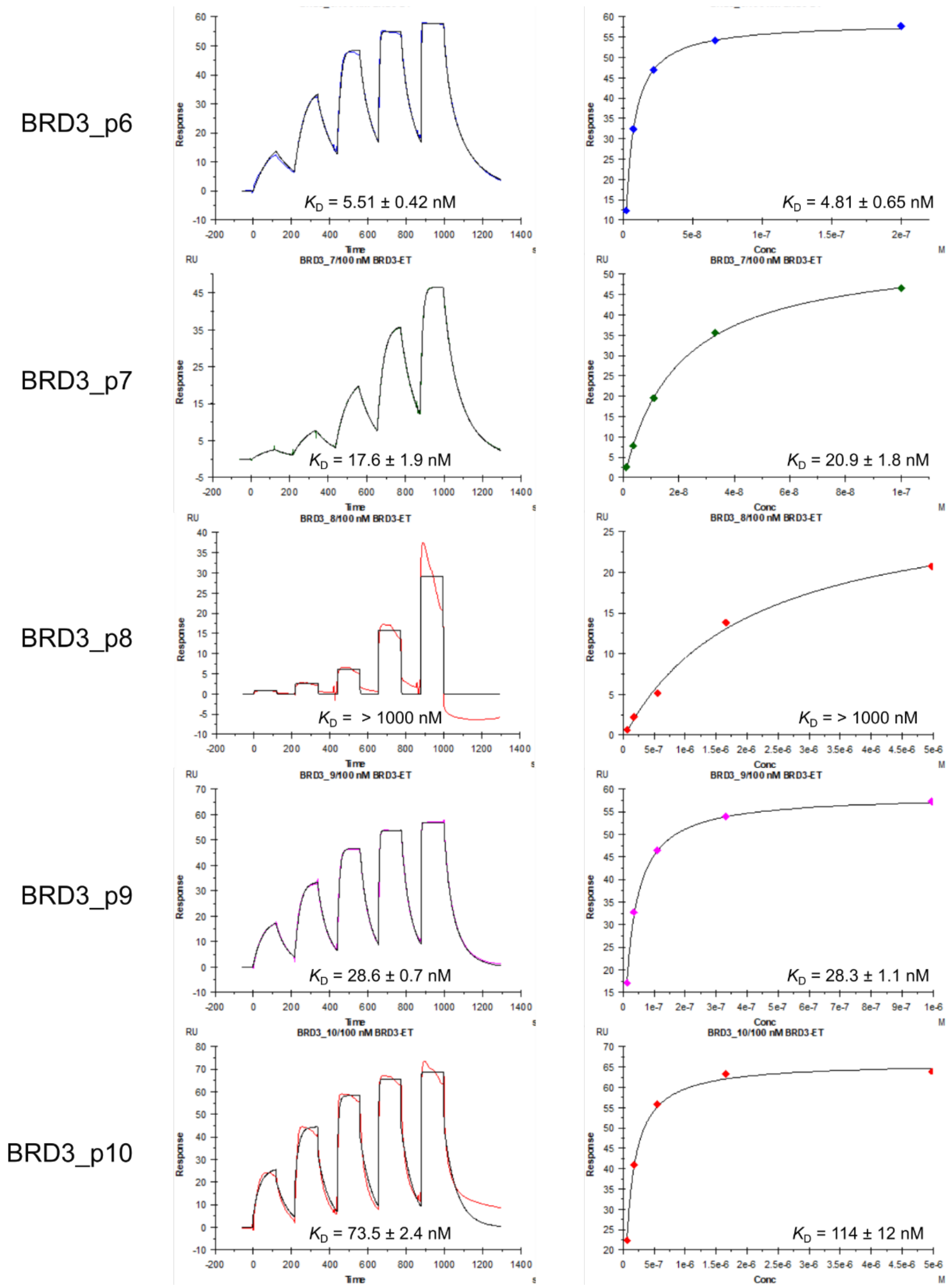

**Figure S9.** Representative SPR data for BRD3-ET. Reported values are the mean $K_D$ ± one standard deviation from two replicates.

**BRD3-L** **Kinetics** **Affinity**

BRD3_p1

BRD3_p2

BRD3_p3.1

BRD3_p3.2

BRD3_p4

BRD3_p5

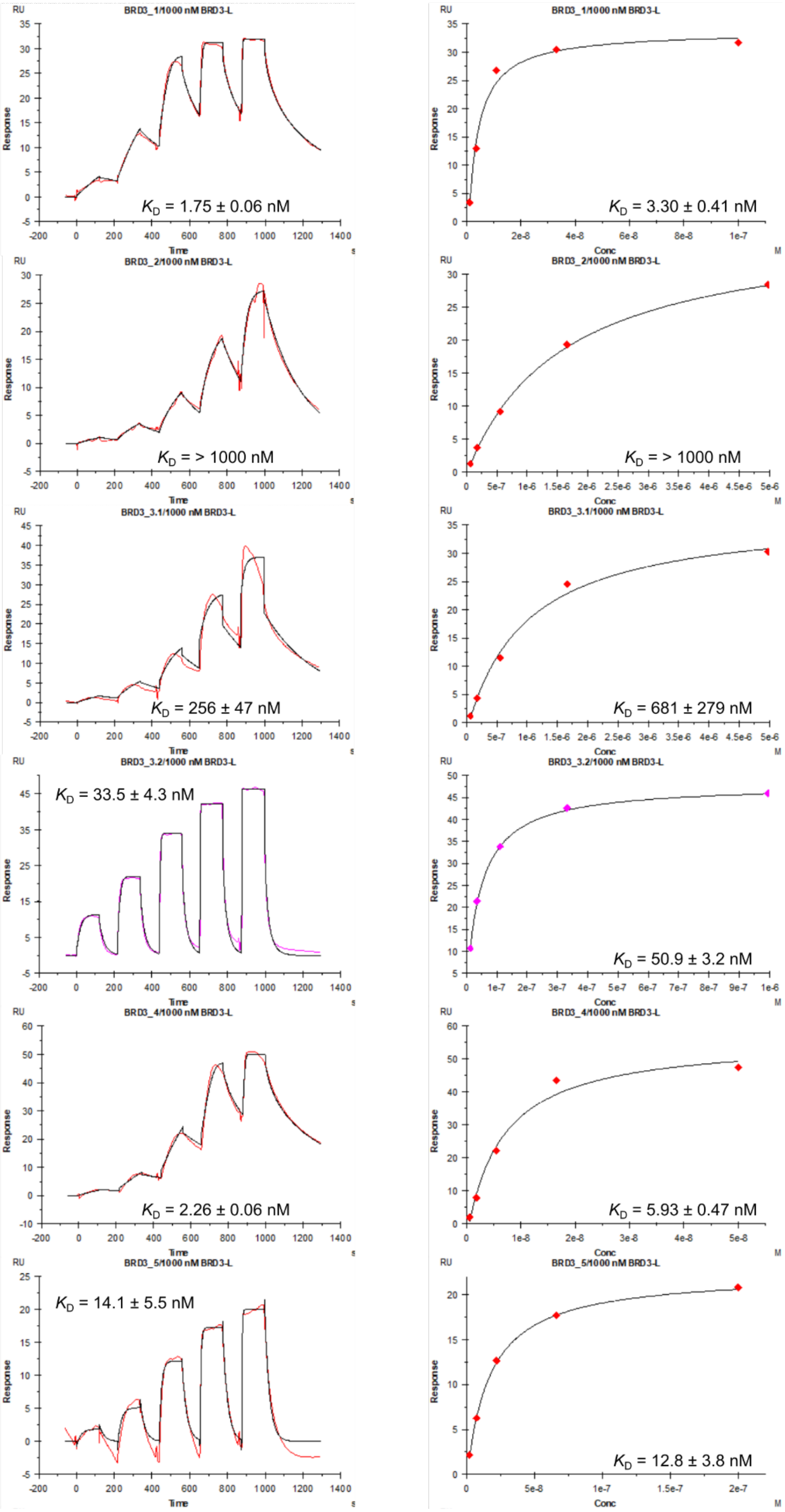

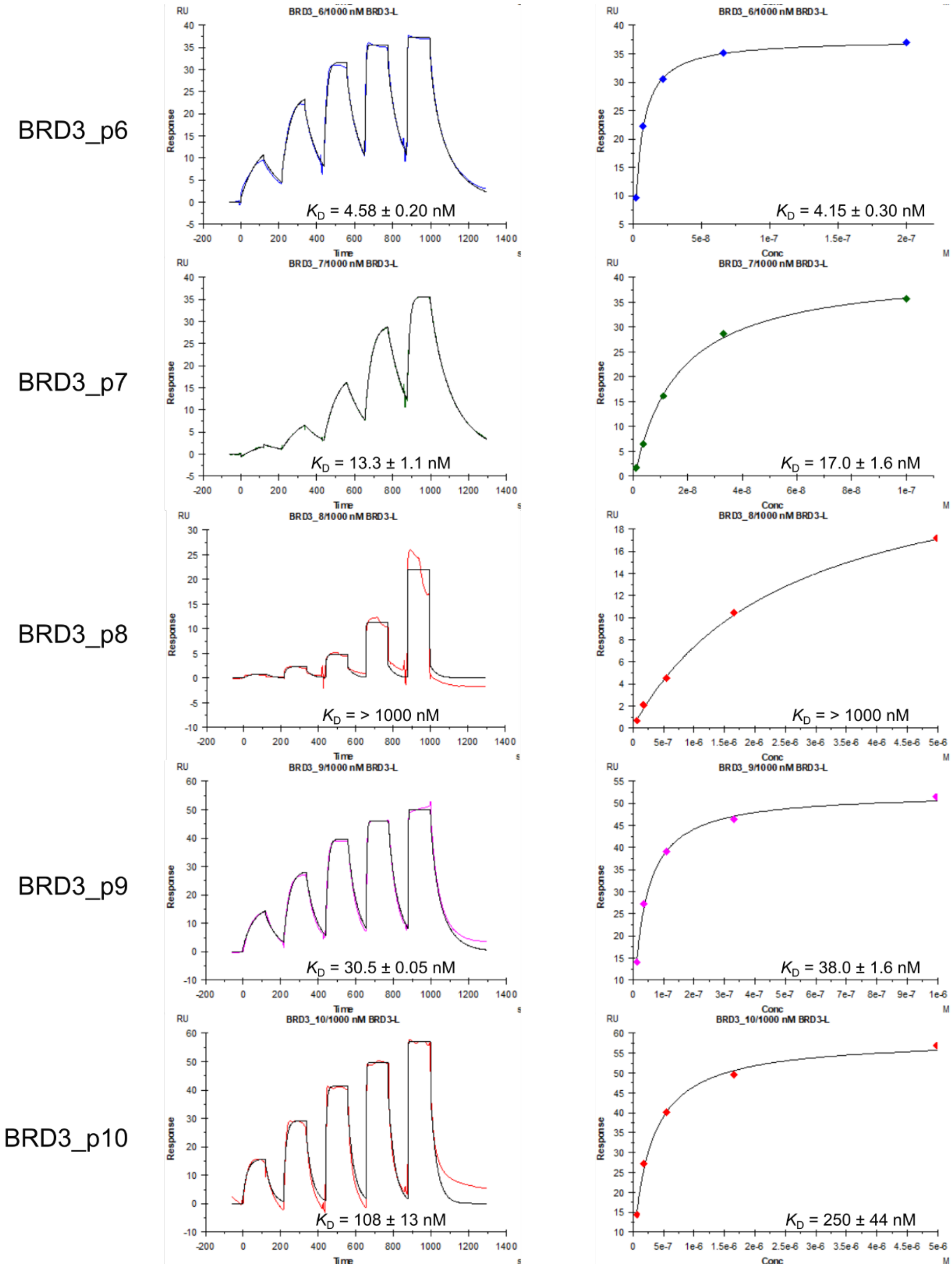

**Figure S10.** Representative SPR data for BRD3-L. Reported values are the mean $K_D$ ± one standard deviation calculated from two replicates.

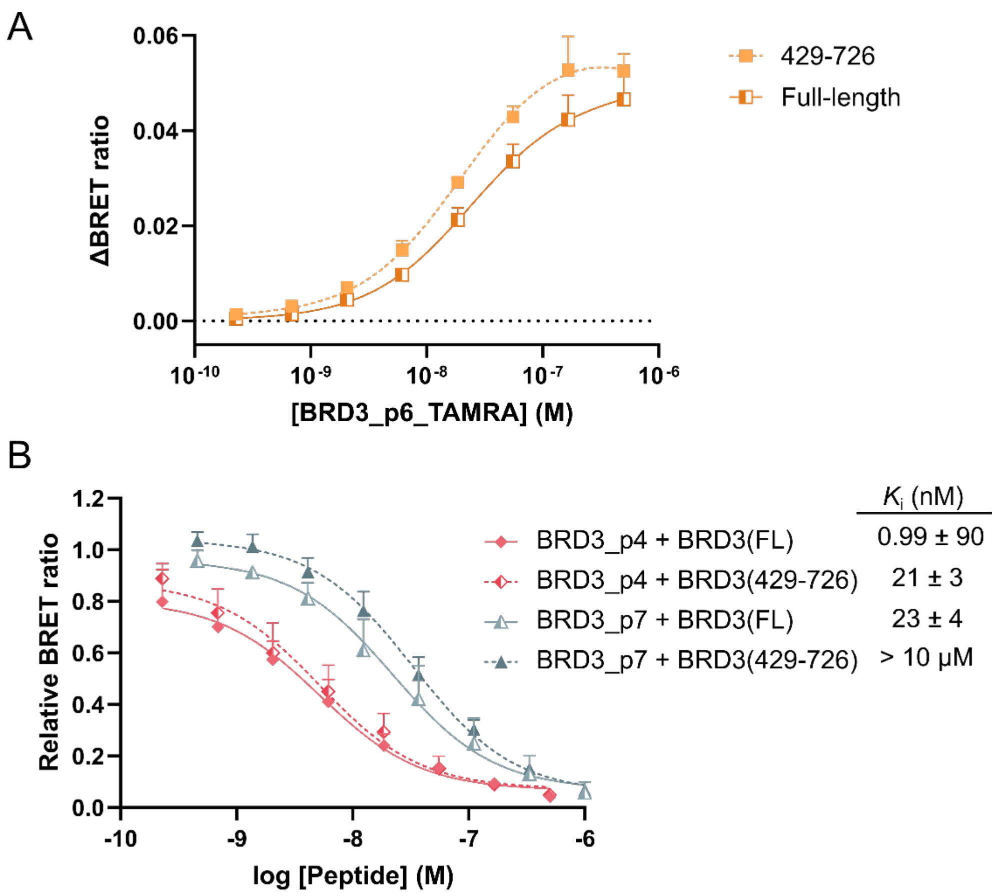

**Figure S11. Peptide binding to BRD3(429-726). A.** Direct binding of BRD3_6_TAMRA to HiBiT-tagged BRD3(Full-length) and BRD3(429-726). **B.** NanoBRET competitions competing BRD3_6_TAMRA (60 nM) from HiBiT-tagged BRD3(429-726). The data for full-length (FL) BRD3 is as shown in Figure 3E. The data displayed are the mean values + one standard error of the mean from at least three independent replicates.

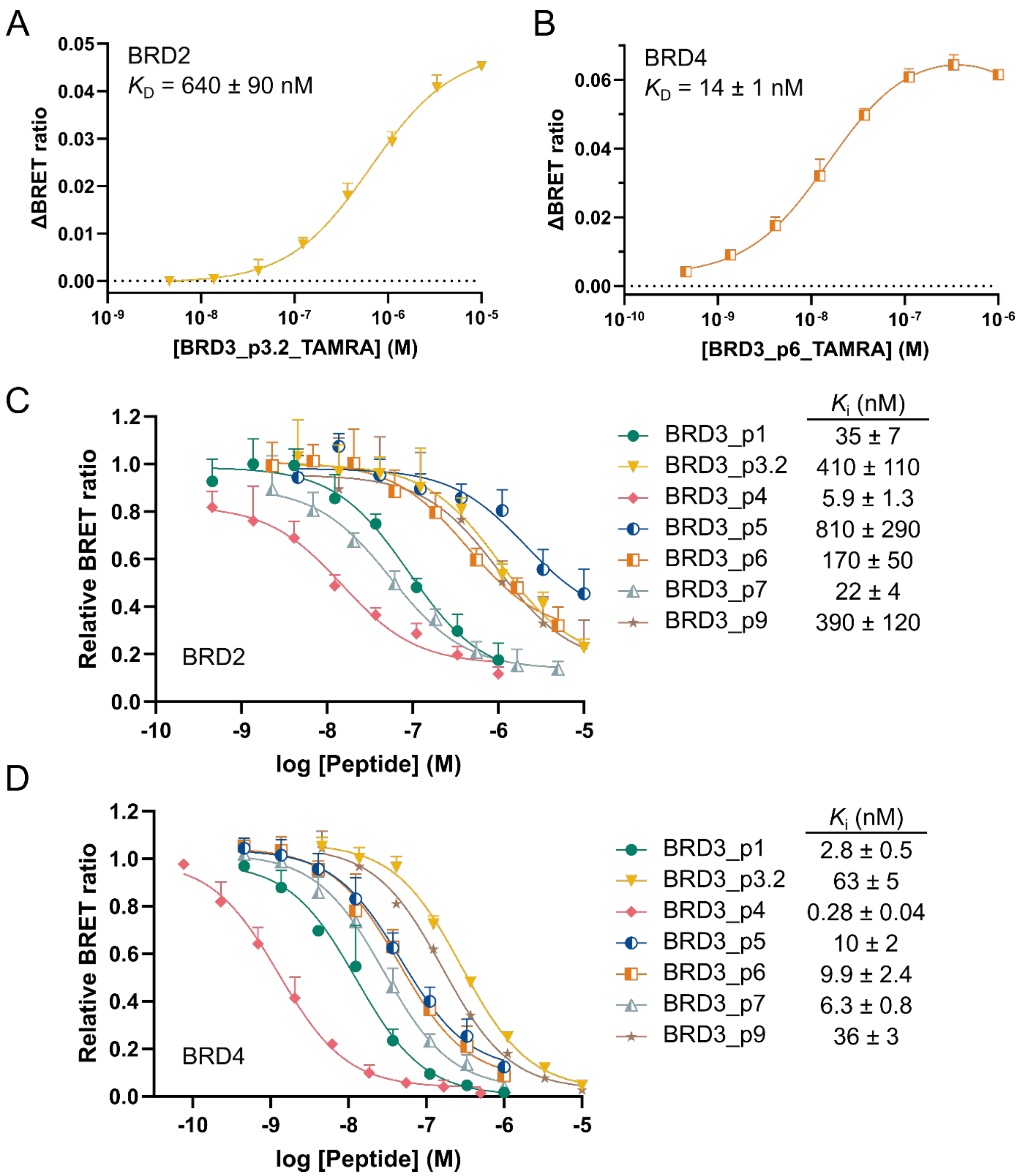

**Figure 12. NanoBRET with full-length BRD2 and BRD4. A.** Direct binding of BRD3_3.2_TAMRA to BRD2-HiBiT. **B.** Direct binding of BRD3_6_TAMRA to BRD4-HiBiT. **C.** Displacement of BRD3_3.2_TAMRA (1 µM) from BRD2-HiBiT by selected peptides in competition assays. **D.** Displacement of BRD3_6_TAMRA (50 nM) from BRD4-HiBiT by selected peptides in competition assays. The data displayed are the mean values + one standard error of the mean from at least three independent replicates.

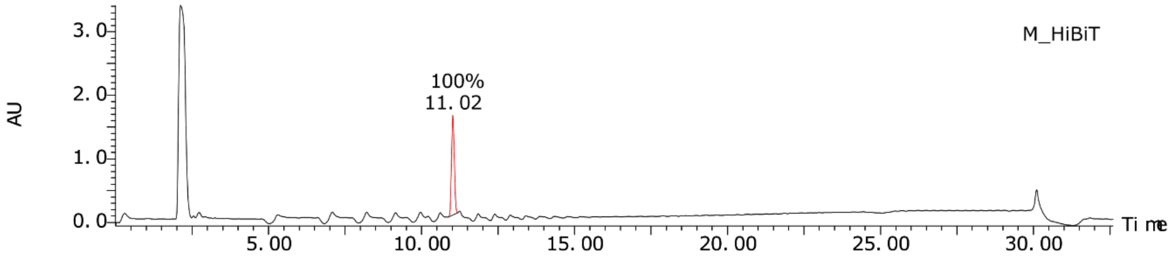
M_HiBiT
AU
3.0
2.0
1.0
0.0
100%
11.02
5.00
10.00
15.00
20.00
25.00
30.00
Time

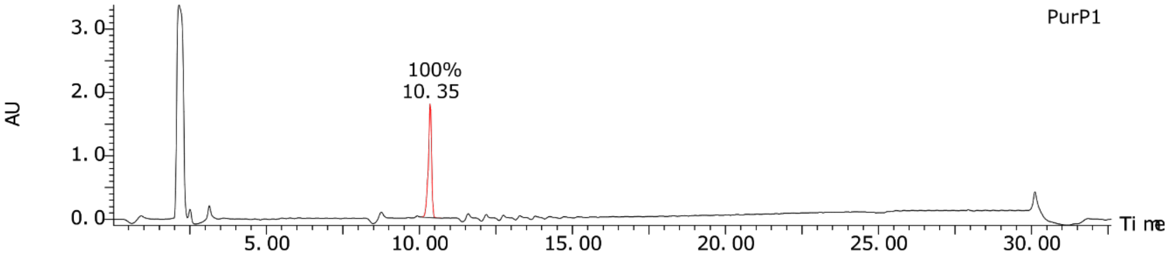
PurP1
AU
3.0
2.0
1.0
0.0
100%
10.35
5.00
10.00
15.00
20.00
25.00
30.00
Time

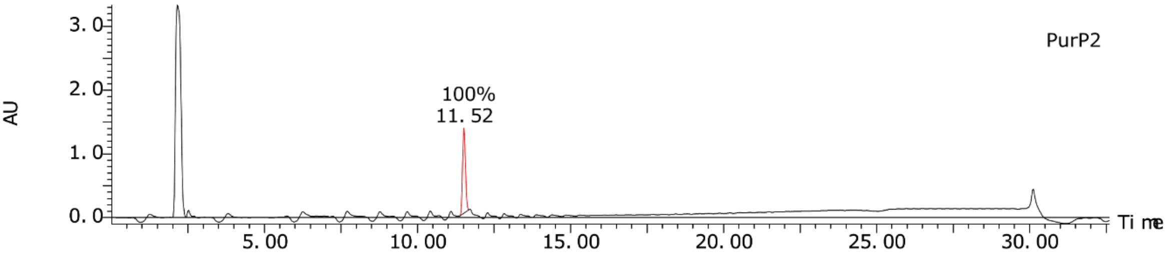
PurP2
AU
3.0
2.0
1.0
0.0
100%
11.52
5.00
10.00
15.00
20.00
25.00
30.00
Time

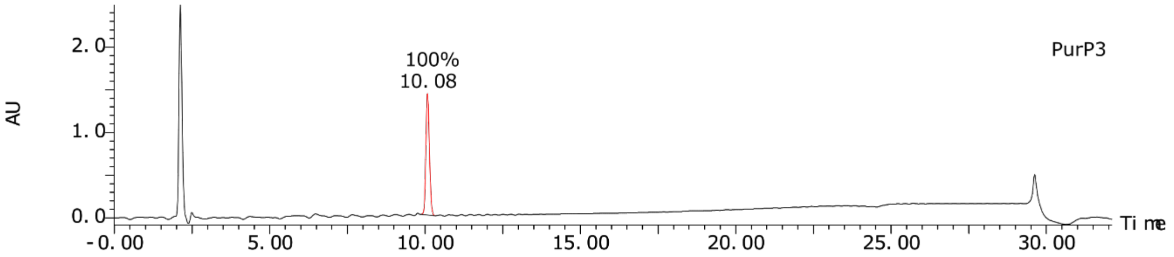
PurP3
AU
2.0
1.0
0.0
100%
10.08
-0.00
5.00
10.00
15.00
20.00
25.00
30.00
Time

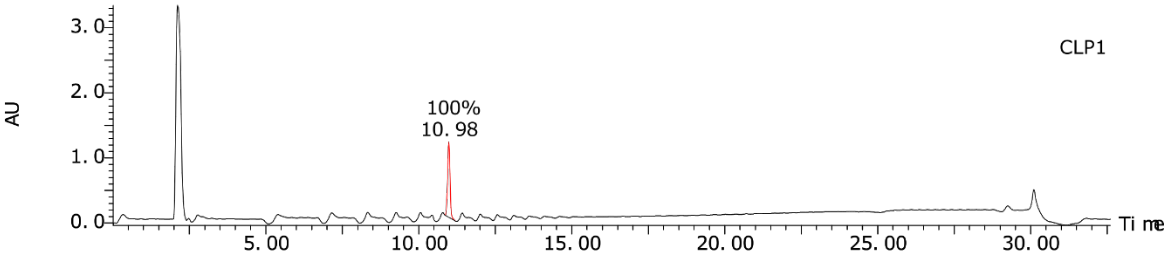
CLP1
AU
3.0
2.0
1.0
0.0
100%
10.98
5.00
10.00
15.00
20.00
25.00
30.00
Time

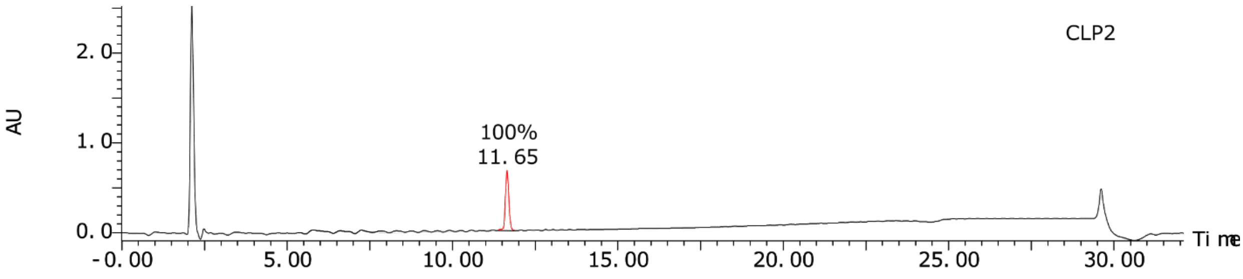
CLP2
AU
2.0
1.0
0.0
100%
11.65
-0.00
5.00
10.00
15.00
20.00
25.00
30.00
Time

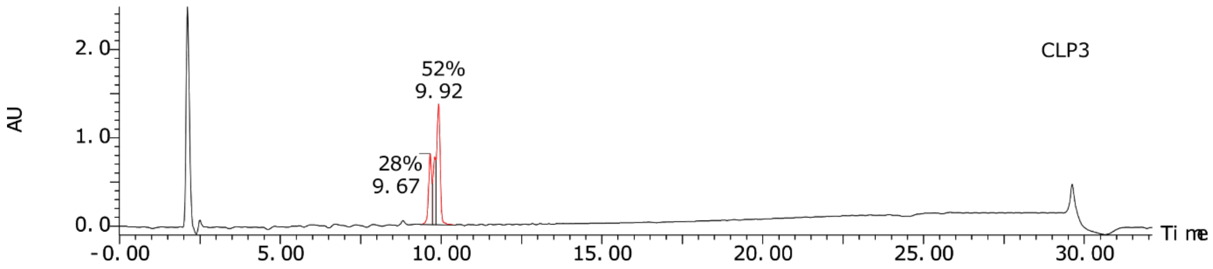

**Figure S13.** Analytical HPLC traces for LgBiT peptides.

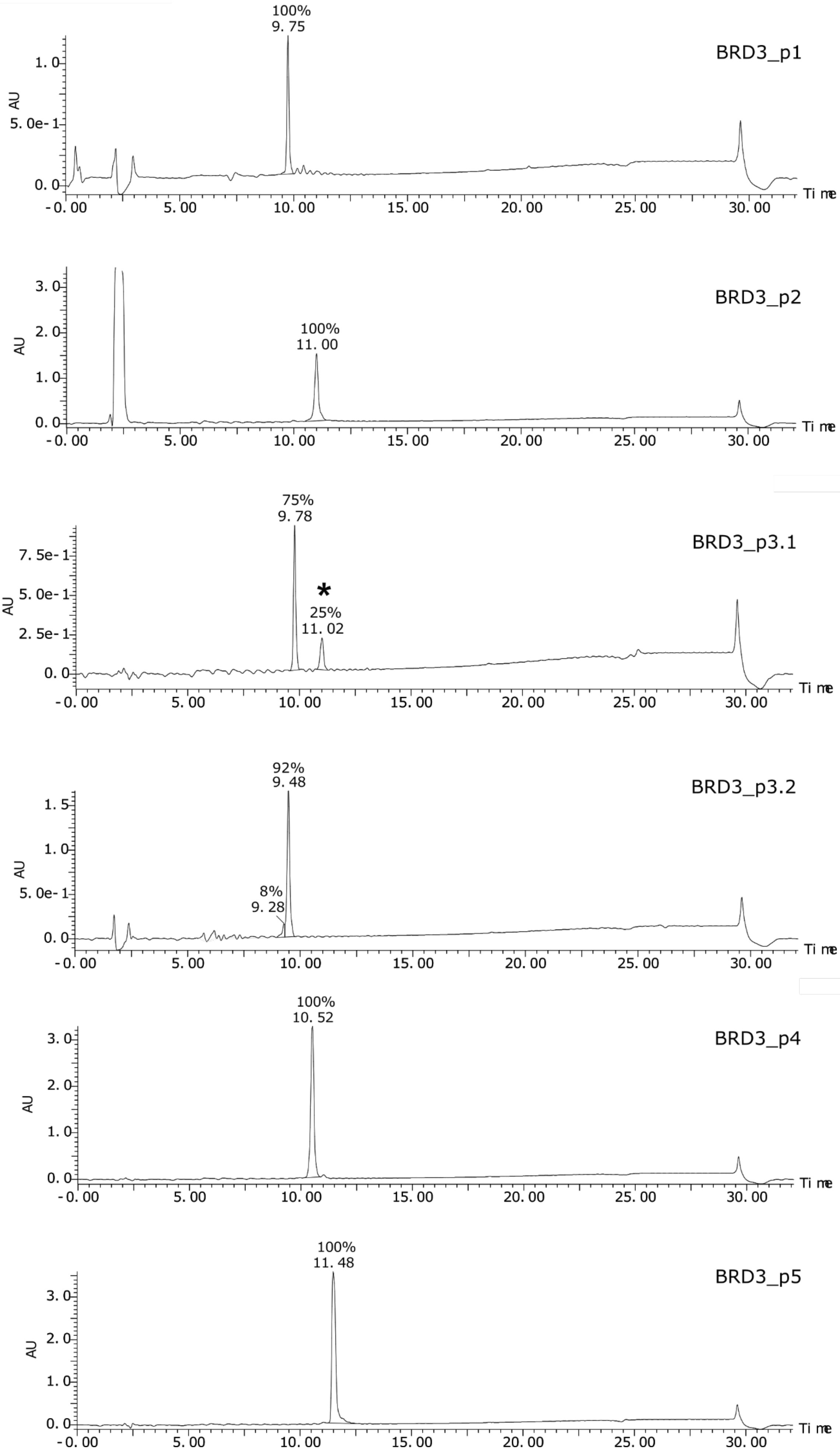
BRD3_p1
100%
9. 75
BRD3_p2
100%
11. 00
BRD3_p3.1
75%
9. 78
*
25%
11. 02
BRD3_p3.2
92%
9. 48
8%
9. 28
BRD3_p4
100%
10. 52
BRD3_p5
100%
11. 48
AU
Time
-0. 00
5. 00
10. 00
15. 00
20. 00
25. 00
30. 00

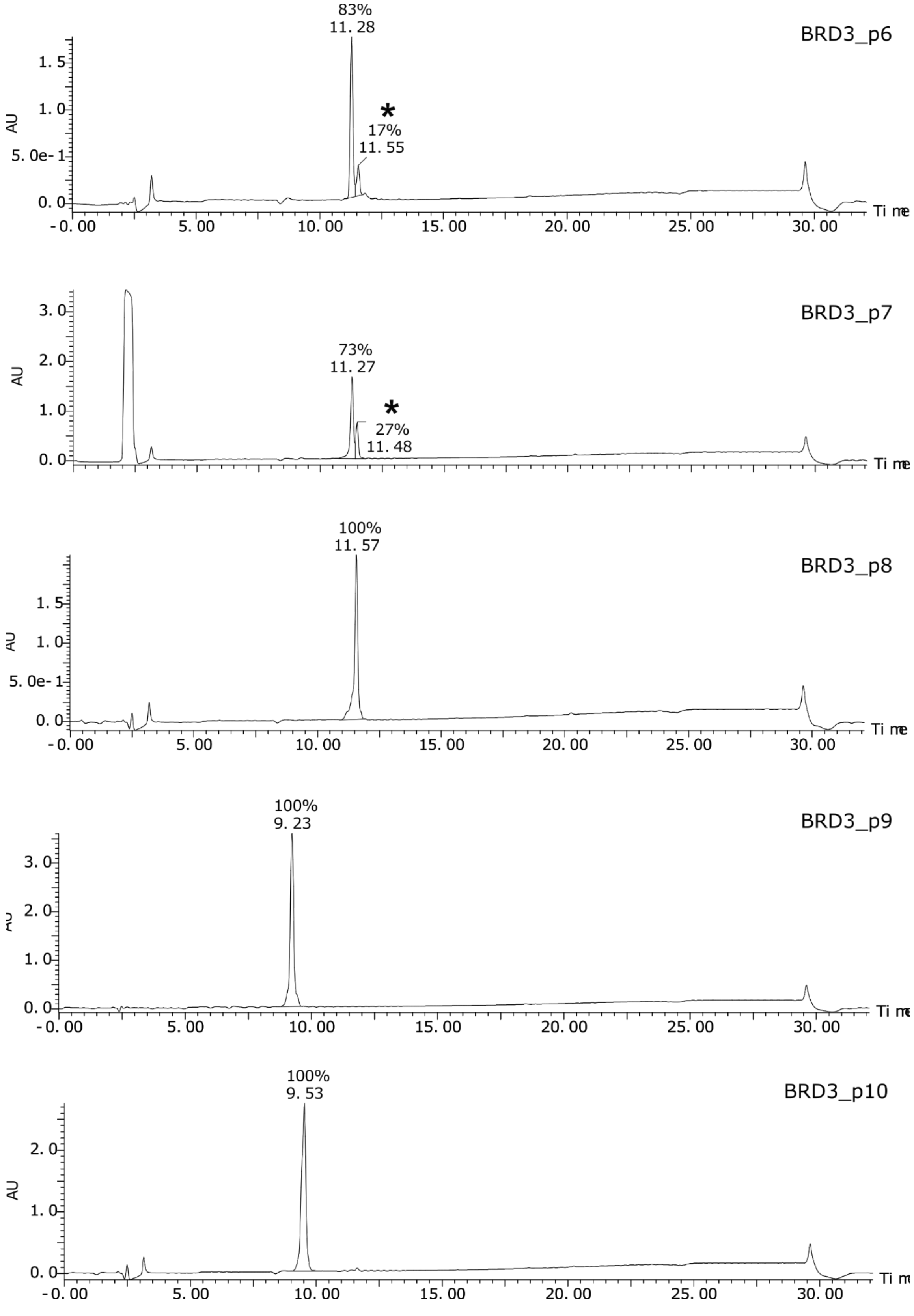

**Figure S14.** Analytical HPLC traces for BRD3 peptides. Peaks marked * are impurities likely resulting from a single amino acid truncation during the synthesis.

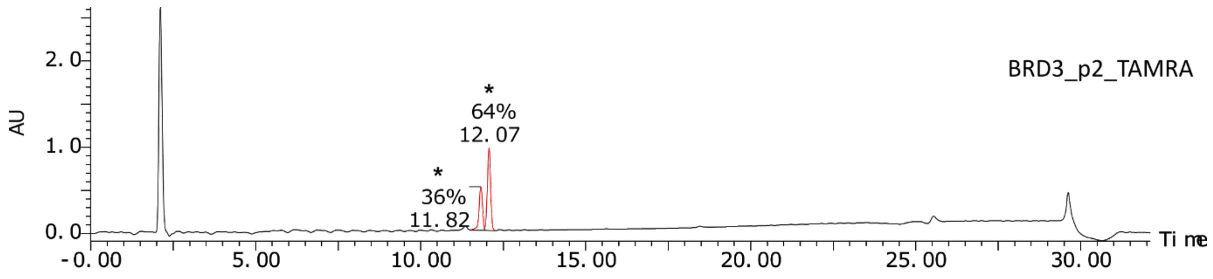
BRD3_p2_TAMRA
AU
2. 0
1. 0
0. 0
*
64%
12. 07
*
36%
11. 82
-0. 00
5. 00
10. 00
15. 00
20. 00
25. 00
30. 00
Time

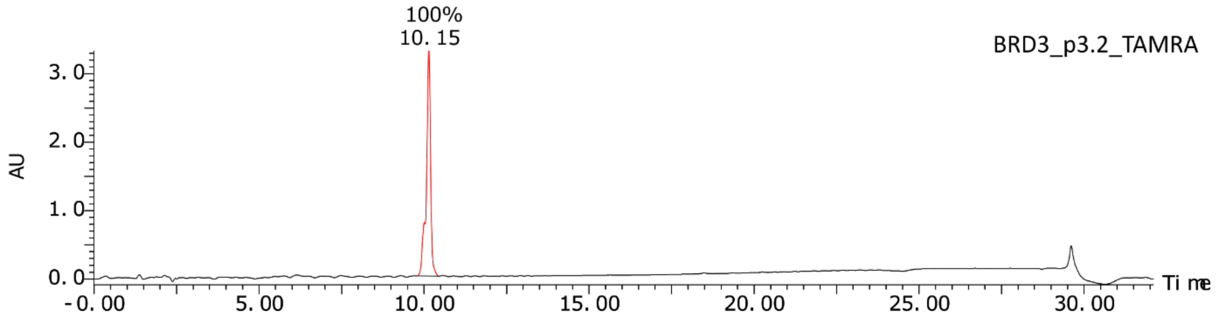
BRD3_p3.2_TAMRA
AU
3. 0
2. 0
1. 0
0. 0
100%
10. 15
-0. 00
5. 00
10. 00
15. 00
20. 00
25. 00
30. 00
Time

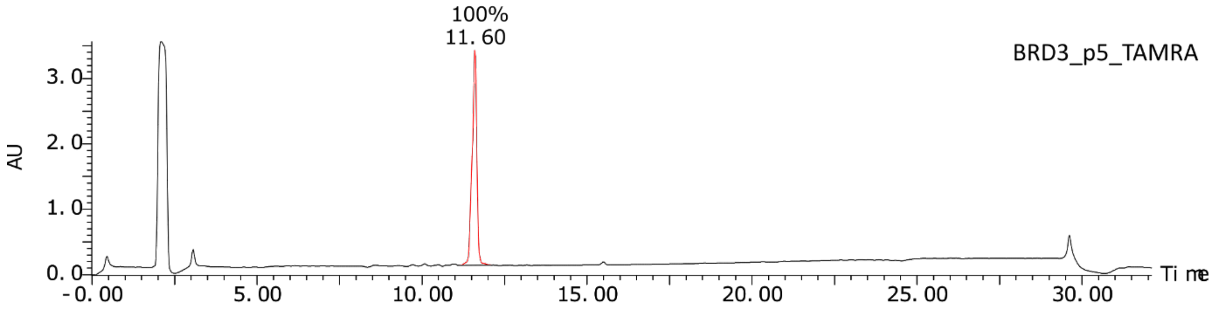
BRD3_p5_TAMRA
AU
3. 0
2. 0
1. 0
0. 0
100%
11. 60
-0. 00
5. 00
10. 00
15. 00
20. 00
25. 00
30. 00
Time

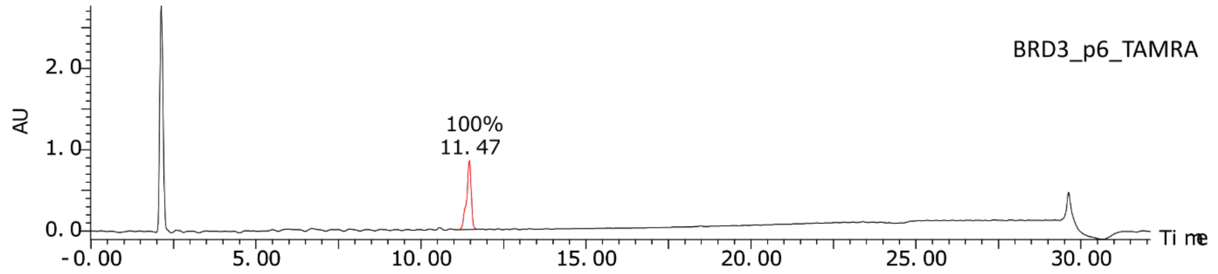
BRD3_p6_TAMRA
AU
2. 0
1. 0
0. 0
100%
11. 47
-0. 00
5. 00
10. 00
15. 00
20. 00
25. 00
30. 00
Time

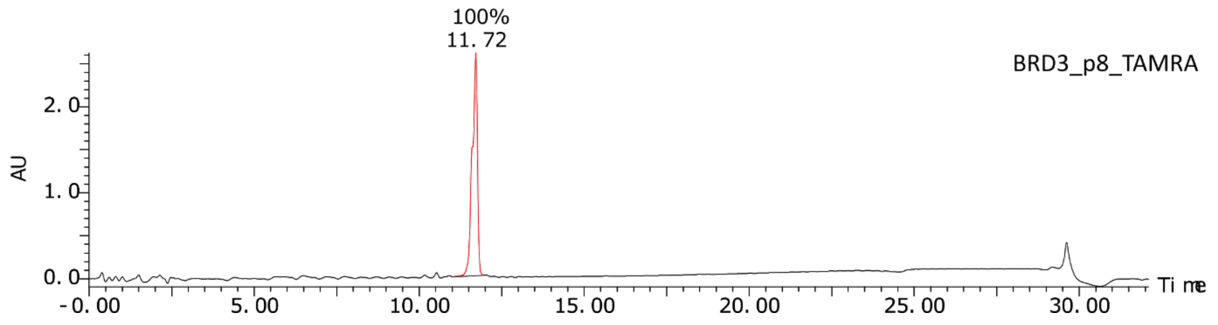
BRD3_p8_TAMRA
AU
2. 0
1. 0
0. 0
100%
11. 72
-0. 00
5. 00
10. 00
15. 00
20. 00
25. 00
30. 00
Time

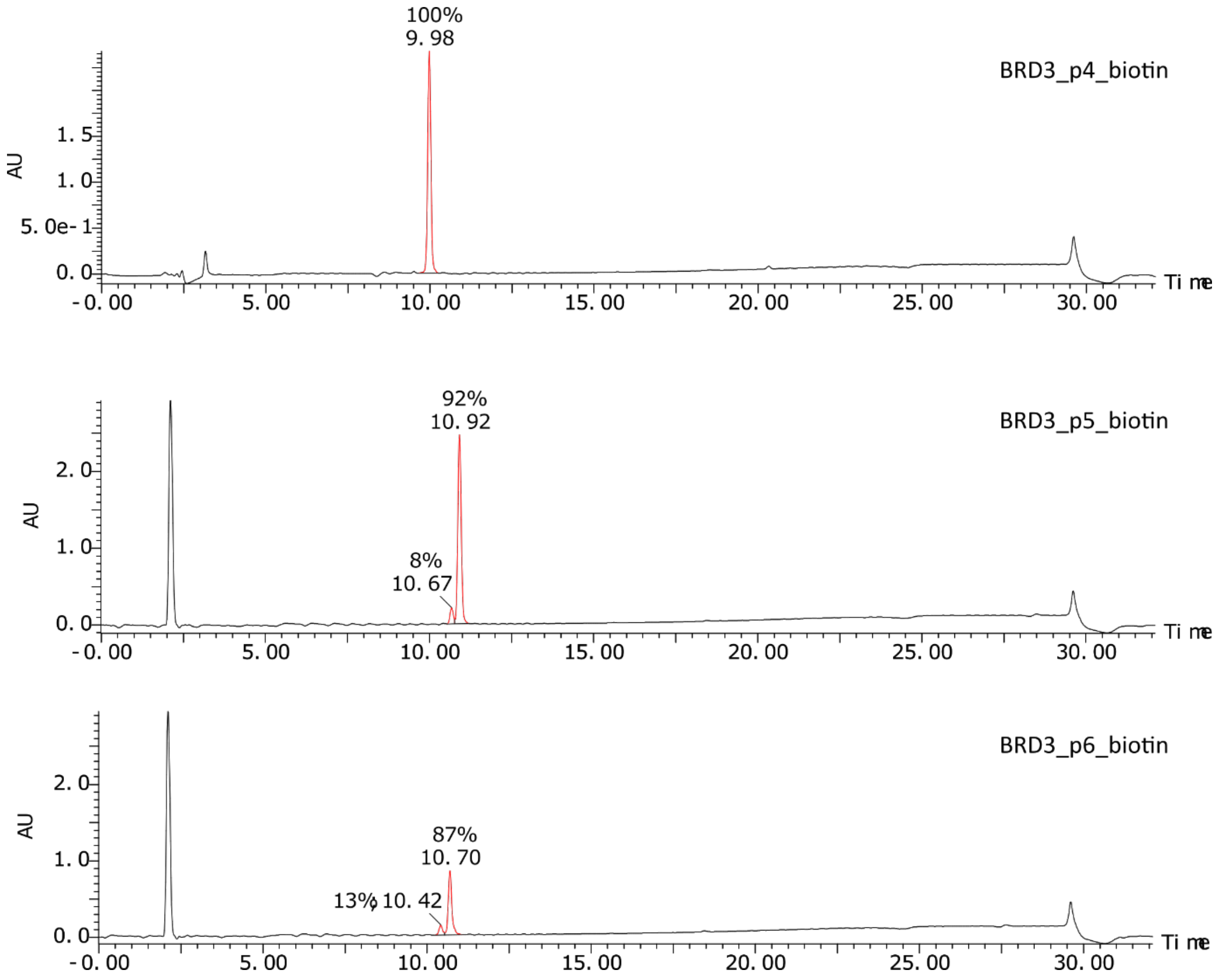

**Figure S15.** Analytical HPLC traces for click-modified BRD3 peptides. For BRD3_2_TAMRA the two peaks (marked *) arise due to the two isomers of TAMRA that are present in the labelling mixture. For other TAMRA-labelled peptides these were mostly separated during purification, however a slight shoulder is visible on the major peak.

**Table S1: LgBiT peptides**

| Name | Sequence | M+H$^+$ | (M+2H$^+$)/2 | Observed |
|---|---|---|---|---|
| M_HiBiT | Ac-MVSGWRLFKKIS-$NH_2$ | 1493.9 | 747.5 | 1493.5/747.5 |
| PurP1 | Ac-MWLRLRRSNKLI-$NH_2$ | 1628.0 | 814.5 | 814.5 |
| PurP2 | Ac-MWIRLLRTNRLN-$NH_2$ | 1628.0 | 814.5 | 1627.8/814.5 |
| PurP3 | Ac-MLFVYKKTQRLN-$NH_2$ | 1583.0 | 792.0 | 1583.0/792.0 |
| CLP1 | Ac-MWILSKRTTRLN-$NH_2$ | 1560.9 | 781.0 | 1560.9/781.1 |
| CLP2 | Ac-MLLILRRTNRLM-$NH_2$ | 1572.0 | 786.5 | 786.6/524.9 |
| CLP3 | Ac-MMSLWKRSNRLN-$NH_2$ | 1577.9 | 789.5 | 1577.7/789.3 |

Ac - acetylated N-terminus

**Table S2: BRD3 peptides**

| Name | Sequence | M+H$^+$ | (M+2H$^+$)/2 | (M+3H$^+$)/3 | Observed |
|---|---|---|---|---|---|
| BRD3_p1 | yRILRIINRNYVVCGS$_\beta$A<u>K</u>-$NH_2$ | 2204.6 | 1102.8 | 735.5 | 736.0 |
| BRD3_p2 | ySPITIINRSVVYCGS$_\beta$A<u>K</u>-$NH_2$ | 2037.3 | 1019.2 | 679.8 | 1019.2 |
| BRD3_p3.1 | yRYLTIVNKVKCISGS $_\beta$A<u>K</u>-$NH_2$ | 2109.5 | 1055.3 | 703.8 | 1055.2/703.3 |
| BRD3_p3.2 | yRYLTIVNKVKSICGS$_\beta$A<u>K</u>-$NH_2$ | 2109.5 | 1055.3 | 703.8 | 1055.1/703.4 |
| BRD3_p4 | yRYITIYWKRYVLCGS$_\beta$A<u>K</u>-$NH_2$ | 2349.8 | 1175.4 | 783.9 | 1175.3 |
| BRD3_p5 | yRYIWIIDRAIVLCGS$_\beta$A<u>K</u>-$NH_2$ | 2206.6 | 1103.8 | 736.2 | 2207.8/1105.0 |
| BRD3_p6 | yQVIWIINQRFVVCGS$_\beta$A<u>K</u>-$NH_2$ | 2190.6 | 1095.8 | 730.9 | 2192.3/1095.3 |
| BRD3_p7 | yRSHWIVLQSVSFCGS$_\beta$A<u>K</u>-$NH_2$ | 2134.8 | 1067.9 | 712.3 | 1067.7/711.8 |
| BRD3_p8 | y**A**YGFILRINRNCGS$_\beta$A<u>K</u>-$NH_2$ | 2026.3 | 1013.7 | 676.1 | 1013.6/675.7 |
| BRD3_p9 | yKLPWTKTHTWHCGS$_\beta$A<u>K</u>-$NH_2$ | 2110.4 | 1055.7 | 704.1 | 1056.1/703.9 |
| BRD3_p10 | yR**A**YHIA**A**LRYKLCGS$_\beta$A<u>K</u>-$NH_2$ | 2178.6 | 1089.8 | 726.9 | 1089.6 |

y = D-Tyr, **A** = *N*-methyl-alanine, $_\beta$A = beta-alanine, <u>**K**</u> = azidolysine

**Table S3: Modified BRD3 peptides**

| Name | Sequence | M+H$^+$ | (M+2H$^+$)/2 | (M+3H$^+$)/3 | Observed |
|---|---|---|---|---|---|
| BRD3_p2_TAMRA | ySPITIINRSVVYCGS$_{\beta}$AK-PEG$_4$-TAMRA | 2683.1 | 1342.1 | 895.0 | 895.2 |
| BRD3_p3.2_TAMRA | yRYLTIVNKVKSICGS$_{\beta}$AK-PEG$_4$-TAMRA | 2755.3 | 1378.1 | 919.1 | 918.7 |
| BRD3_p5_TAMRA | yRYIWIIDRAIVLCGS$_{\beta}$AK-PEG$_4$-TAMRA | 2852.4 | 1426.7 | 951.5 | 951.3 |
| BRD3_p6_TAMRA | yQVIWIINQRFVVCGS$_{\beta}$AK-PEG$_4$-TAMRA | 2836.3 | 1418.7 | 946.1 | 945.7 |
| BRD3_p8_TAMRA | yAYGFILRINRNCGS$_{\beta}$AK-PEG$_4$-TAMRA | 2672.1 | 1336.6 | 891.4 | 890.8 |
| BRD3_p4_biotin | yRYITIYWKRYVLCGS$_{\beta}$AK-PEG$_4$-biotin | 2807.4 | 1404.2 | 936.5 | 1405.2/936.9 |
| BRD3_p5_biotin | yRYIWIIDRAIVLCGS$_{\beta}$AK-PEG$_4$-biotin | 2664.2 | 1332.6 | 888.7 | 2663.6/1333.7 |
| BRD3_p6_biotin | yQVIWIINQRFVVCGS$_{\beta}$AK-PEG$_4$-biotin | 2648.2 | 1324.6 | 883.4 | 2648.2/1325.4 |

y = D-Tyr, **A** = *N*-methyl-alanine, $_{\beta}$A = beta-alanine, **K** = azidolysine

**Table S4. Primers used for mRNA display**

| Oligo name | Sequence |
|---|---|
| T7g10M.F46 | TAATACGACTCACTATAGGGTTAACTTTAAGAAGGAGATATACATA |
| CGS3an13.R39 | TTTCCGCCCCCCGTCCTAGCTGCCGCTGCCGCTGCCGCA |
| GS3an13.R36 | TTTCCGCCCCCCGTCCTAGCTGCCGCTGCCGCTGCC |
| CGS3an13.R22 | TTTCCGCCCCCCGTCCTAGCTG |
| M_HiBiT.R75 | GCTGCCGCTGCCGCTGCCAGAGATTTTCTTGAACAGACGCCAACCAGAAACCAT ATGTATATCTCCTTCTTAAAG |
| BRD3.1b.F74 | AAGAAGGAGATATACATATGAAAACCATTATGGGCATGACCTGGCGCACCATG CAGTGCGGCAGCGGCAGCGGC |
| 11aa_linear.R75 | GCTGCCGCTGCCGCTGCCMNNMNNMNNMNNMNNMNNMNNMNNMNNM NNMNNCATATGTATATCTCCTTCTTAAAG |
